# Convergent evolution of the army ant syndrome and congruence in big-data phylogenetics

**DOI:** 10.1101/134064

**Authors:** Marek L. Borowiec

## Abstract

The evolution of the suite of morphological and behavioral adaptations underlying the ecological success of army ants has been the subject of considerable debate. This “army ant syndrome” has been argued to have arisen once or multiple times within the ant subfamily Dorylinae. To address this question I generated data from 2,166 loci and a comprehensive taxon sampling for a phylogenetic investigation. Most analyses show strong support for convergent evolution of the army ant syndrome in the Old and New World but certain relationships are sensitive to analytics. I examine the signal present in this data set and find that conflict is diminished when only loci less likely to violate common phylogenetic model assumptions are considered. I also provide a temporal and spatial context for doryline evolution with time-calibrated, biogeographic, and diversification rate shift analyses. This study underscores the need for cautious analysis of phylogenomic data and calls for more efficient algorithms employing better-fitting models of molecular evolution.

**Significance:** Recent interpretation of army ant evolution holds that army ant behavior and morphology originated only once within the subfamily Dorylinae. An inspection of phylogenetic signal in a large new data set shows that support for this hypothesis may be driven by bias present in the data. Convergent evolution of the army ant syndrome is consistently supported when sequences violating assumptions of a commonly used model of sequence evolution are excluded from the analysis. This hypothesis also fits with a simple scenario of doryline biogeography. These results highlight the importance of careful evaluation of signal and conflict within phylogenomic data sets, even when taxon sampling is comprehensive.

## Introduction

Army ants (Figure 1) are a charismatic group of organisms that inspire research in such disparate fields as behavioral ecology (Schöning et al., 2005), biodiversity conservation (Peters et al., 2008), and computational biology (Garnier et al., 2013). These ants are distributed throughout warm temperate and tropical regions of the world and belong to a more inclusive clade known as the subfamily Dorylinae (Brady et al., 2014). They are characterized by a suite of morphological and behavioral adaptations, together dubbed the army ant syndrome (Brady, 2003). This syndrome includes obligate collective foraging, frequent colony relocation, and highly specialized wingless queens. In contrast to many other ant species, army ants never forage individually. Nests of army ants are temporary and in some species colonies undergo cycles of stationary and nomadic phases (Schneirla, 1945). Unable to fly, the queens must disperse on foot and are adapted to producing enormous quantities of brood (Raignier et al., 1955). Other peculiarities of army ants include colony reproduction by fission and highly derived male morphology. No army ant species are known to lack any of the components of the syndrome (Gotwald, 1995). Because of its antiquity and persistence, the army ant syndrome has been cited as an example of remarkable long-term evolutionary stasis (Brady, 2003). Several distantly related lineages of ants evolved one or more of the components of the army ant syndrome but did not reach the degree of social complexity or ecological dominance of the “true army ants” in the Dorylinae (Kronauer, 2009).

**Figure 1:**
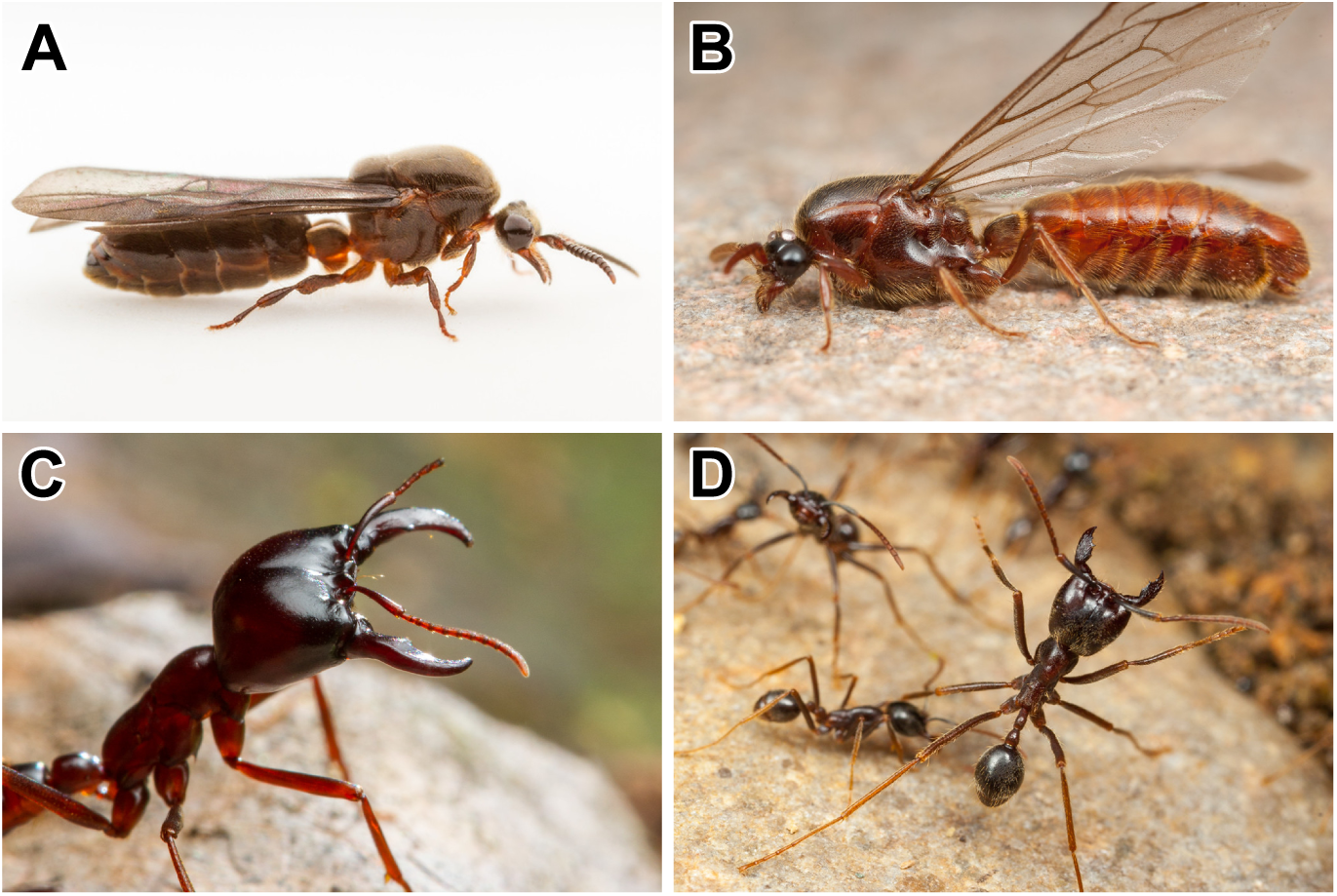
Striking morphological similarity of Old World (left) and New World (right) army ants. A: *Aenictus* male, B: *Neivamyrmex* male, C: *Dorylus* soldier, D: *Labidus* soldier. Photographs courtesy of Alex Wild/alexanderwild.com.

A major question of army ant biology is whether the army ant syndrome originated only once or if it arose independently in New World and Old World army ants (Kronauer, 2009). The answer requires a robust phylogenetic framework, but Dorylinae are an example of an ancient rapid radiation and elucidating its phylogeny has proven difficult (Brady et al., 2014). Before the advent of quantitative phylogenetic methods, the army ants were thought to have arisen more than once within the subfamily (Brown, 1975; Gotwald, 1979; Bolton, 1990). This view was based on the observation that army ants are poor at dispersal and on the assumption that they evolved after the breakup of Gondwana around 100 Ma ago. More recent studies (Baroni Urbani et al., 1992; Brady, 2003), however, reported monophyly of army ant lineages, even though statistical support for this grouping was often low (Brady et al., 2014). Furthermore, the most recent divergence time estimates suggest that army ants indeed originated after the breakup of Gondwana, implying either independent origins or long-distance dispersal (Kronauer, 2009).

Here I reassess the phylogeny of the Dorylinae including New World and Old World army ants. I use a large phylogenomic data set and taxon sampling increased twofold compared to a previous phylogeny (Brady et al., 2014), including 155 taxa representing more than 22% of described doryline species diversity and all 27 currently recognized extant genera (Borowiec, 2016b). The sequence data comes from a total of 2,166 loci centered around Ultraconserved Elements or UCEs (Faircloth et al., 2012; Branstetter et al., 2017) distributed throughout the ant genomes, comprising 892,761 nucleotide sites with only 15% of missing data and gaps.

Analyses of the complete data set and loci with high average bootstrap support or “high phylogenetic signal” (Salichos and Rokas, 2013) produce results that are sensitive to partitioning scheme and inference method. Analysis of data less prone to systematic bias, such as slow-evolving (Rodríguez-Ezpeleta et al., 2007; Betancur-R. et al., 2013; Goremykin et al., 2015) or compositionally homogeneous loci (Jermiin et al., 2004) and amino acid sequences (Hasegawa and Hashimoto, 1993), consistently support the hypothesis of independent origins of the army ant syndrome in the Old and New World.

## Results

### Relationships Among Doryline Lineages Inferred From Combined Data Matrix

The concatenated dataset produces different topologies depending on the partitioning scheme and method used to infer the phylogeny (Figure 2A–D, P). The true army ants are not monophyletic in the maximum likelihood tree under partitioning by locus. The tree shows a clade that includes almost all New World dorylines, including New World army ants (hereafter called “New World Clade”; Figure 2A; Supplementary Figure 1). The latter include the genera *Cheliomyrmex*, *Eciton*, *Labidus*, *Neivamyrmex*, and *Nomamyrmex*. Apart from New World army ants, the New World Clade unites all exclusively New World lineages of the Dorylinae and includes *Acanthostichus*, *Cylindromyrmex*, *Leptanilloides*, *Neocerapachys*, and *Sphinctomyrmex*. Old World driver ants *Dorylus* and the poorly known genus *Aenictogiton* are sister to *Aenictus*. Other well-supported clades comprising multiple genera include a clade of South-East Asian and Malagasy lineages *Cerapachys*, *Chrysapace*, and *Yunodorylus*, a well-resolved clade of several Old World genera (*Lioponera*, *Lividopone*, *Parasyscia*, *Zasphinctus*) that was also recovered in (Brady et al., 2014). Another South-East Asian clade consists of the mostly subterranean genera *Eusphinctus*, *Ooceraea*, and *Syscia*. The backbone of the tree shows multiple short internodes and generally lower resolution, consistent with a previous study (Brady et al., 2014). The maximum likelihood tree inferred from the same data set but using k-means partitioning (Frandsen et al., 2015) shows a strongly-supported Old World *Aenictus* and New World army ants clade (Figure 2B; Supplementary Figure 2). Army ants are not monophyletic in this tree either, because *Dorylus* and *Aenictogiton* are not sister to the *Aenictus* plus New World army ants clade. Unpartitioned analysis results in a similar topology (Figure 2C; Supplementary Figure 3).

**Figure 2:**
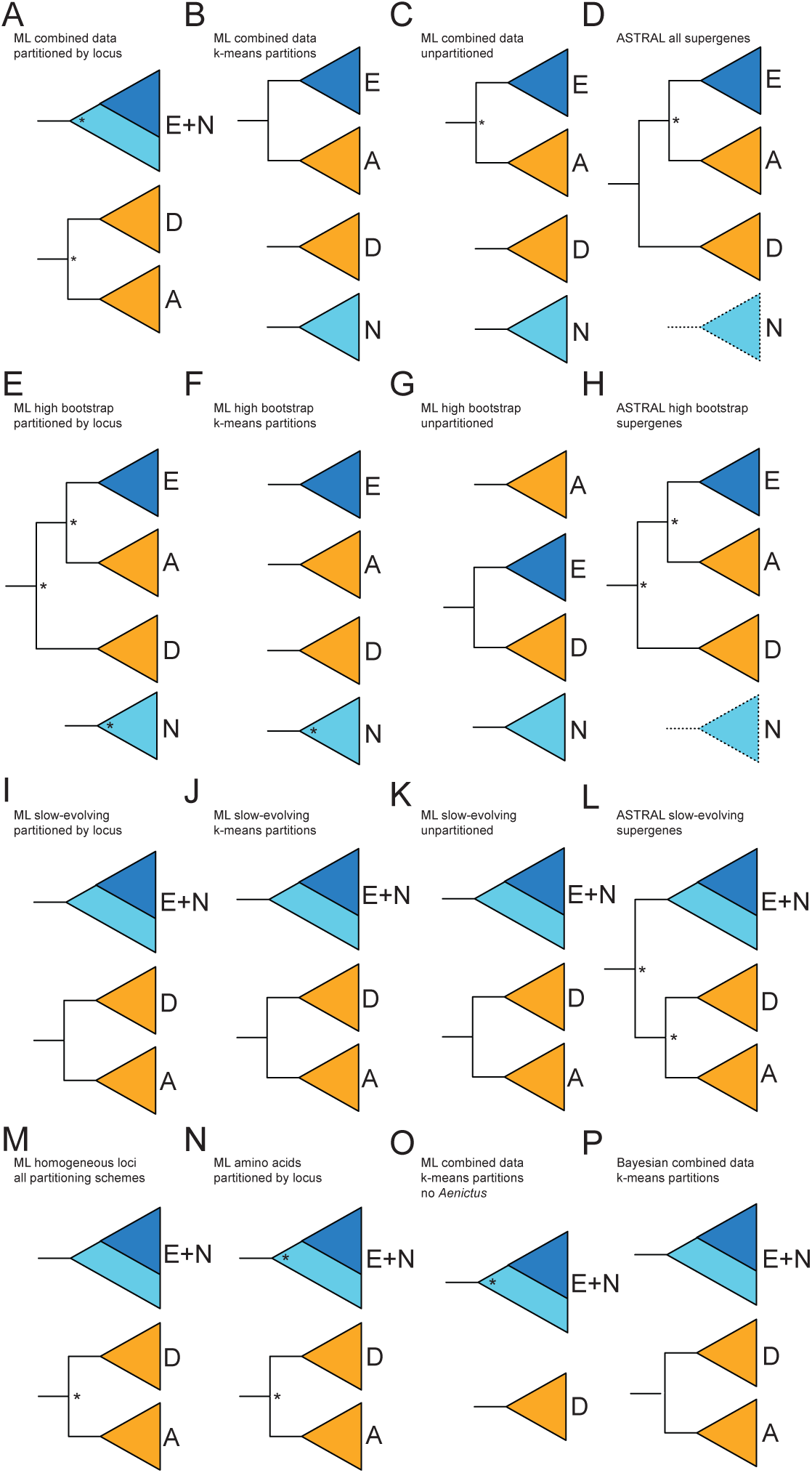
Summary of phylogenetic position of army ant lineages recovered in different analyses. In each figure, letter A signifies *Aenictus*, D: *Aenictogiton*+*Dorylus*, E: New World army ants (*Cheliomytmex*, *Eciton*, *Labidus*, *Neivamyrmex*, *Nomamyrmex*), N: all other New World dorylines (*Acanthostichus*, *Cylindromyrmex*, *Leptanilloides*, *Neocerapachys*, *Sphinctomyrmex*) except *Syscia*. Note increased incongruence in analysis of combined data set (A–D) and “high signal” data set (E–H) compared to slow-evolving (I–L) and compositionally homogeneous (M) loci. Asterisk indicates support *≤* 99%. Dashed lines signify polyphyly. See Supplementary Figures 1–18 for complete trees.

Coalescent species tree inferred from all loci is similar to the concatenation results with respect to poor resolution at deep nodes (Figure 2D; Supplementary Figure 4). The species tree resembles the unpartitioned and k-means partitioned concatenated trees because of moderate support for the *Aenictus* plus New World army ants clade, and contrast most with both concatenated trees by supporting army ant monophyly, the hypothesis favored by previous studies (Brady, 2003; Brady and Ward, 2005; Brady et al., 2014). Overall support for the deep nodes is lower in species trees and this approach does not recover some of the clades present across all concatenated analyses, namely the New World or South-East Asian/Malagasy groups mentioned above.

### Evidence for Bias in Maximum Likelihood Tree Based on All and “High Signal” Loci

The conflict among ML topologies derived from different partitioning schemes and between concatenated and species tree warranted further investigation. I constructed five data sets in addition to the combined data matrix: matrix of “high signal” loci (Salichos and Rokas, 2013) equivalent in length to 1/5 of the sites in combined data set matrix, matrix composed of slow-evolving loci equivalent in length to 1/5 of the sites in combined data set matrix, matrix composed of only compositionally homogeneous loci, and matrix that excluded the long-branched *Aenictus* species but is otherwise identical to combined data matrix (See Supplementary Table 2 for more information on these matrices). Additionally, I developed a workflow to extract putatively protein-coding regions from the UCE data set (see Extended Methods) and analyzed those regions separately, coded as amino acids.

Analyses of loci whose trees have highest mean bootstrap support (high phylogenetic signal sensu Salichos and Rokas (2013)) shows even more discordance among analyses than that of the combined data set (Figure 2E–H; Supplementary Figures 5–8). Army ant lineages show different topologies in each of the three used partitioning schemes.

Using only the slowest evolving one-fifth of the data (”slow-evolving” hereafter), there is universal support for a clade that unites all exclusively New World lineages, including the New World army ants under all partitioning schemes (Figure 2I–K; Supplementary Figures 9–11). Species tree analyses of slow-evolving loci also support New World Clade and the *Aenictus* plus (*Aenictogiton*+*Dorylus*) clade (Figure 2L; Supplementary Figure 12).

Examining locus properties in different rate bins reveals that more rapidly evolving sequences and “high signal” loci exhibit qualities previously associated with systematic bias, namely higher compositional heterogeneity (Lockhart et al., 1994; Jermiin et al., 2004) and saturation (Philippe and Forterre, 1999). The most slowly evolving loci have lower overall among-taxon sequence heterogeneity than more rapidly evolving and high bootstrap loci (Supplementary Figure 28A; RCFV two-sample t-test slow-evolving vs. all other loci: *t* = 28.564, slow-evolving *mean* = 0.0358, all other loci *mean* = 0.0575, *p* « 0.01; slow-evolving vs. high bootstrap: *t* = 27.538 high bootstrap *mean* = 0.0647, *p* « 0.01). Saturation is also greater in more rapidly evolving and high bootstrap loci ((Supplementary Figure 28B; slope of regression two-sample t-test slow-evolving vs. all other loci: *t* = 21.682, slow-evolving *mean* = 0.474, all other loci *mean* = 0.361, *p* « 0.01; slowevolving vs. high bootstrap: *t* = 10.129, high bootstrap *mean* = 0.393, *p* « 0.01). Of the 379 loci that pass the homogeneity test (Foster, 2004), 244 are found among slow-evolving loci and zero are present among the high bootstrap loci.

Analysis of loci that pass the phylogeny-corrected compositional homogeneity test (Foster, 2004) shows universal support for the New World Clade, regardless of partitioning scheme (Figure 2M; Supplementary Figure 13–15). Among-taxon compositional heterogeneity is a relatively well-understood source of bias in phylogenetic analysis (Jermiin et al., 2004) and is not accounted for by the general time-reversible model of sequence evolution used in RAxML, the program used here for maximum likelihood inference.

Amino acid alignments are also known to be more robust against compositional bias (Hasegawa and Hashimoto, 1993) and saturation, although not free from it when distantly related lineages are considered (Foster and Hickey, 1999). In this study the amino acid matrix of protein-coding sequences is highly conserved compared to the nucleotide matrix of combined data with 14% proportion of parsimony informative sites in the former compared to 48% in the latter. The amino acid matrix analyzed under maximum likelihood recovers monophyletic New World Clade and Old World army ants (Figure 2N; Supplementary Figure 16).

More evidence for the artefactual nature of the grouping of Old World and New World army ants comes from an analysis where *Aenictus* is removed from the combined data matrix partitioned under the k-means scheme. If *Aenictus* was not affecting the position of New World army ants on the tree, one would expect to see no change of the position of the latter if the former is removed. This is not the case here; the phylogeny recovered from the alignment without *Aenictus* has New World army ants nested within the New World Clade (Figure 2O; Supplementary Figure 17).

Bayesian analysis of k-means partitioned combined data matrix strongly supports the New World Clade and monophyly of *Aenictogiton*+*Dorylus*, unlike the ML analysis of the same dataset under the same partitioning scheme and sequence evolution model (Figure 2P; Supplementary Figure 18).

In summary, although concatenated ML and species tree analysis of combined data matrix and high-bootstrap loci support grouping of Old World army ants *Aenictus* with New World army ants under some conditions, analyses using more reliable data are congruent in their support for independent origins of the army ant syndrome in New World and Old World.

### The Timeline of Doryline Evolution and Diversification

Fossilized birth-death (FBD) process divergence dating (Heath et al., 2014) employed here shows that crown doryline ants started diversifying in the Cretaceous, around 74 Ma (53–101 Ma 95% highest probability density interval or HPD) ago according to Bayesian inference (97 Ma under penalized likelihood or PL; Supplementary Figure 19) (Figure 3). The FBD results near the root are characterized by high uncertainty but the mean age suggests a younger crown age than 87 Ma recovered in a previous study (Brady et al., 2014). The difference is likely at least in part due to a different calibration approach used in the present study (FBD versus node dating) and a revised, younger age estimate of Baltic amber (Aleksandrova and Zaporozhets, 2008). Similar to concatenation, divergence dating shows a tree highly compressed during early evolution, with 16 splits occurring within the first 20 Ma. According to penalized likelihood this period of early diversification lasted longer, accounting for about 35 Ma. The most recent common ancestor of the New World Clade is resolved at 60 Ma (43–83 Ma 95% HPD interval; 60 Ma under PL). The split of Old World army ant lineages of *Aenictus* and *Dorylus*+*Aenictogiton* is apparently very ancient at 58 Ma (41–79 Ma 95% HPD; 54 Ma under PL) and occurred during the initial diversification period. The old age of this node and long branch subtending extant *Aenictus* help explain why this relationship is difficult to recover. The five currently recognized genera of New World army ants share an ancestor at about 28 Ma (20–37 Ma 95% HPD; 27 Ma under PL) ago. These dates are younger than those inferred for several of the non-army ant doryline genera such as *Eburopone* or *Leptanilloides*. The conspicuous above-ground foraging seen in some *Dorylus* driver ants and in New World *Eciton* is remarkably young, as it appears that these groups diversified within the last 5–6 Ma, a result robust across different dating analyses (see Supplementary Figures 19-21). The driver ant genus *Dorylus* is shown to be young relative to earlier analyses (Brady, 2003; Kronauer et al., 2007) at ca. 16 Ma (11–22 95% HPD; 15 Ma under PL).

**Figure 3:**
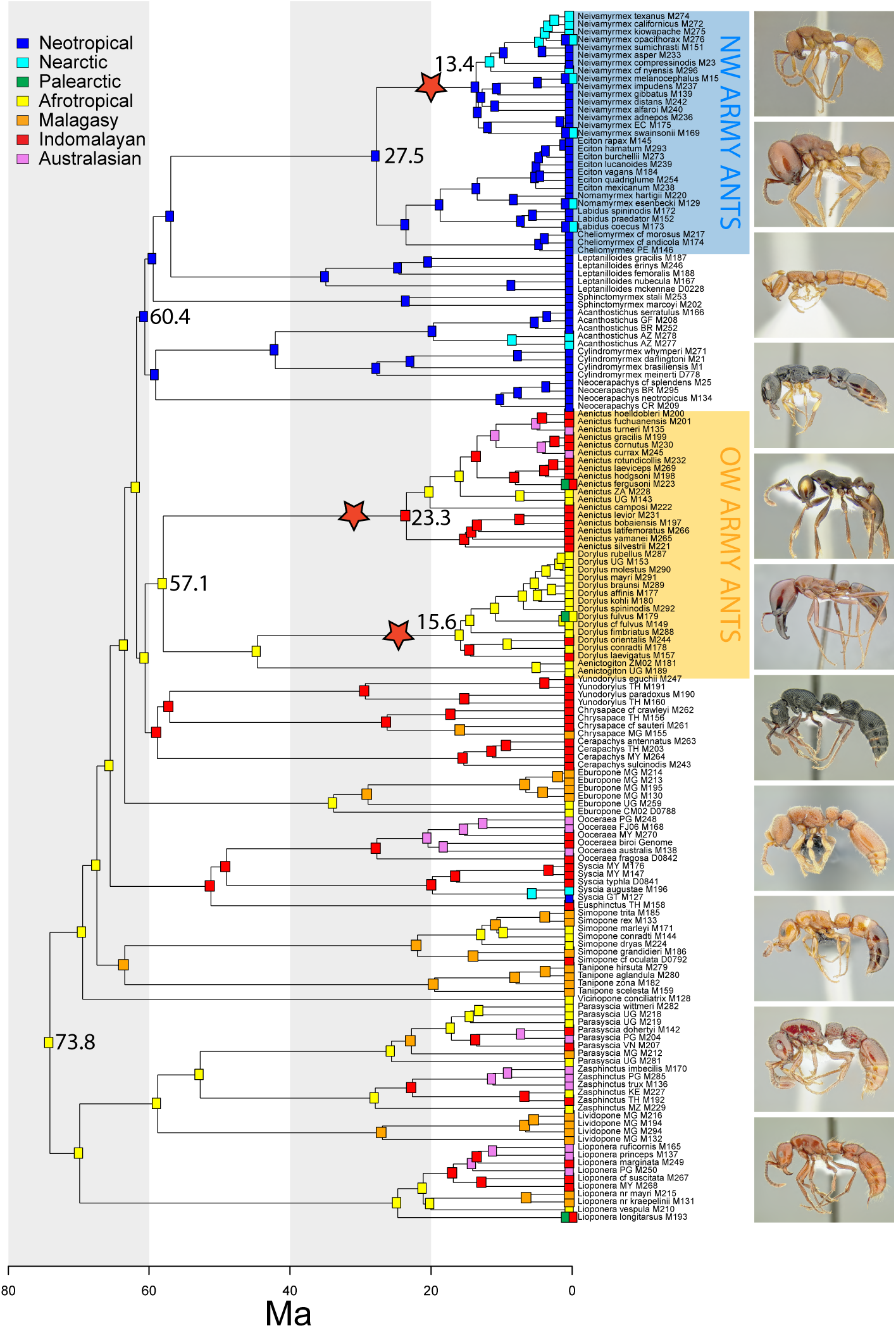
Timeline of doryline evolution and biogeographic history. The highest relative probabilities of ancestral ranges are shown as inferred using BioGeoBEARS under DEC+J, averaged over 100 trees from the BEAST posterior. The tree is the BEAST consensus. Selected mean divergence time estimates are given at nodes. Stars signify rate shifts. All dates in Ma. See Supplementary Figures 20, 21, and 26 for average node dates, posterior probabilities from the BEAST analysis, and pie charts of relative probabilities of all possible ancestral ranges, respectively.

The Bayesian divergence time estimates, especially those early in the tree and outside the New World Clade, are associated with considerable uncertainty. This is likely due to both topological uncertainty and the fact that only seven fossil calibrations were available for the dorylines, all except one located within the New World Clade (Supplementary Table 4; Supplementary Figure 20).

Diversification shift analyses in BAMM (Rabosky, 2014) identified a three shift-scenario for dorylines, with shifts occurring separately on the branches subtending *Aenictus*, *Dorylus*, and *Neivamyrmex* (Figure 3). Rate shift configurations in which only two shifts were identified, however, were also common in the posterior sampling (Supplementary Figure 25), either on branches leading to *Aenictus* and *Neivamyrmex* only, or on the branches subtending the clade of *Aenictus*, *Aenictogiton*, and *Dorylus* and *Neivamyrmex* (see Supplementary Figures 22-25).

### Biogeographic History

Strong geographic affinities are apparent within the doryline phylogeny. Large clades are mostly confined to only one or two adjacent realms, although movement within both the Old and New World appears to have been common (Figure 3). There is strong evidence for Old World origins of the dorylines and the analyses summarized across a sample of trees from the posterior suggest an Afrotropical ancestral range (Figure 3; Supplementary Figure 26), although the analysis under BEAST consensus only results in high uncertainty and combined Afrotropical-Malagasy-Oriental as the most likely ancestral range (Supplementary Figure 27). Two lineages are confined to the New World. One gave rise to the radiation of almost all extant New World forms, including New World army ants and their kin. The dates estimated for the origin of this New World Clade coincide with warm climatic conditions and multiple land bridges connecting the Old and New World in northern latitudes (Brikiatis, 2014). The other New World group is much younger and appeared to arrive some time after 28 Ma ago. The presumably SE Asian or Palearctic ancestor of the New World species of *Syscia* either crossed the Beringian land bridge or dispersed across the Pacific further south, since by that time North Atlantic connections were closed (Sanmartín et al., 2001). The Old World *Aenictus* and New World genus *Neivamyrmex*, although superficially similar in appearance and biology, illustrate different scenarios of biogeographic history for generic lineages within Dorylinae. Crown *Aenictus* is older at 23 Ma (34 Ma under PL) and originated in the Indomalayan region. It then subsequently moved into the Afrotropics, dispersed back into Indomalaya, and moved into Australasia with possible movement back into Indomalaya. Some species also range into the Palearctic. In contrast, *Neivamyrmex* remained largely confined to the Neotropics where it originated around 13 Ma (20 Ma under PL) ago, with at least one clade moving into and diversifying in the southern Nearctic and with some species returning to the Neotropics or straddling the boundary of the two adjacent regions. Madagascar is a center of doryline diversity with seven overall and two endemic genera but no true army ants (Figure 3).

### Concluding Remarks

#### Doryline Biology and Evolution Need Further Study

The new phylogeny presented here reveals that the army ant syndrome can be viewed as both an example of long-term evolutionary stasis and a remarkable case of convergent evolution. Brady (2003) argued that the army ant syndrome originated only once around 100 Ma ago and has since persisted without loss in any descendant lineages. While the present study suggests that this set of behavioral and morphological traits evolved at least twice in the Dorylinae, it also shows that the syndrome has been conserved within Old and New World army ants. The alternative scenario of single origin requires multiple losses on lineages leading to both Old and New World army ants (Figure 3), an unlikely proposition given that no species are known to have lost any of the syndrome components in the large and diverse genera such as *Aenictus*, *Dorylus*, or *Neivamyrmex*.

Despite the improved resolution of the army ant tree, much work remains to be done with the regard to doryline evolution. A particularly vexing matter is poor knowledge of the Afrotropical genus *Aenictogiton*. Although based on male morphology and its phylogenetic affinity to *Dorylus* it has been assumed that it is a subterranean army ant, there is no direct evidence of army ant behavior or queen morphology (Borowiec, 2016b). If *Aenictogiton* is not an army ant, our views on the evolution of the army ant syndrome have to be adjusted. In general, the current knowledge of doryline ecology and behavior is mostly limited to the minority of species that are conspicuous above-ground foragers, although the clonal *Ooceraea biroi* is a notable exception (Tsuji and Yamauchi, 1995; Ravary and Jaisson, 2002; Oxley et al., 2014). Better understanding of doryline behavior and morphology will undoubtedly yield further insights into the evolution of the army ant syndrome. For example, independent evolution of army ants is perhaps less surprising given that certain components of the syndrome (e.g., derived queen and male morphologies) appear multiple times within the subfamily. Unfortunately, too little data exists for a rigorous study of these trends in a comparative framework (Borowiec, 2016b). Early anatomical research implied army ant polyphyly by emphasizing differences in sting and Dufour gland morphologies (Hermann, 1969; Billen, 1985; Billen and Gotwald, 1988) between the Old and New World army ants. The new phylogenetic and taxonomic (Borowiec, 2016b) framework should reinvigorate comparative work on doryline morphology and biology.

#### Densely Sampled Phylogenomic Data Sets Are Not Robust to Artefacts and Bias

Genome-scale data offers powerful tools for reconstructing phylogenies. This study, however, demonstrates that caution is necessary when evaluating hypotheses generated by these new data sets, even when taxon sampling is comprehensive. Empirical studies that emphasize the importance of bias often recommend improving taxon sampling but usually also deal with cases where sampling could be increased relative to first attempts, such as in broad-scale phylogenies of eukaryotes or Metazoa (Delsuc et al., 2005). The doryline phylogeny represents a case where the scope for improvement by additional sampling of currently known lineages is limited. This is because adding more species is likely to fail at significantly shortening long branches in the tree (Borowiec, 2016b). Researchers should be thus wary of systematic bias whenever combination of very short and very long branches is encountered, regardless of taxon sampling. Two general strategies are available for exploring sensitivity to model mis-specifications: using only data less prone to bias or applying better-fitting heterogeneous models (Rodríguez-Ezpeleta et al., 2007). The latter approach is ideal but computationally expensive, becoming prohibitive with data sets such as the one used here, including over 150 taxa and thousands of loci.

In this study, reduction of phylogenetic noise resulting from compositional heterogeneity and saturation increased congruence among topologies obtained using different analytics. In the case of the complete data set, mutually exclusive and strongly supported results were be obtained depending on the statistical framework (*e.g.*, maximum likelihood vs. Bayesian) or sequence evolution model (*e.g.*, k-means partitioning vs. partitioning by locus) chosen for analysis. Using only “high signal” loci (Salichos and Rokas, 2013) with highest average bootstrap exacerbated incongruence among analyses. These loci also exhibited undesirable properties such as higher potential for saturation and violation of the among-taxon compositional heterogeneity. This indicates that using additional measures of data quality is needed in phylogenomics, such as direct and indirect measures of model mis-specification (Brown, 2014). Other recent research suggests that analysis results can be strongly affected by a tiny proportion of highly biased loci or sites (Shen et al., 2017). In conclusion, phylogenomic studies should always perform sensitivity analyses to test the robustness of the result to different analytics.

## Acknowledgments

I would like to thank Phil Ward for guidance, support, and valuable conversations related to this project. Many thanks to Michael Branstetter who allowed me to work with an unpublished probe set and provided wet lab protocol training. Alex Wild, Corrie Moreau, and Daniel Kronauer gave important feedback during early stages of conceiving this study. Alex also provided photographs of live army ants used in Figure 1. Brian Johnson and Christian Rabeling kindly gave access to computing resources. Karl Kjer and Kimiora Ward provided comments that helped to improve this manuscript. Thank you to everyone who contributed specimens for sequencing or helped with field work: Fidèle Bemaeva, Brendon Boudinot, Júlio Chaul, Katsuyuki Eguchi, Flávia Esteves, Rodrigo Feitosa, Brian Fisher, Paco Hita-Garcia, Milan Janda, Jack Longino, Andrea Lucky, Sean McKenzie, Matt Prebus, Andry Rakotomalala, Caspar Schöning, Michael Staab, Andy Suarez, and Phil Ward. This research was funded by NSF Doctoral Dissertation Improvement Grant 1402432, a Young Explorers Grant from National Geographic, Microsoft Azure Research Award, Henry A. Jastro awards from the Department of Entomology and Nematology at UC Davis, and facilitated by Ernst Mayr Travel Grants in Animal Systematics and SYNTHESYS awards FR-TAF-594 and GB-TAF-303.

## Methods

### Taxon Sampling and Data Generation

For a detailed description of methods and additional references see Supplementary Materials. I chose doryline species for the analyses based on a recent generic revision of the subfamily (Borowiec, 2016b) and other recent taxonomic and phylogenetic work. I maximized the breadth of sampling by including at least one representative from each biogeographic region in which a genus occurs and aiming to sample across morphologically disparate groups within genera. I extracted the genomic DNA, prepared libraries and enriched them with molecular probes designed as described in (Faircloth et al., 2014), targeting 2,524 loci (Branstetter et al., 2017). Dual-indexed (Faircloth and Glenn, 2012), enriched sequences were sequenced on two lanes of Illumina HiSeq 2500 platform. I cleaned demultiplexed reads using Illumiprocessor (Faircloth, 2011) and performed assembly with Trinity (Grabherr et al., 2011). I then mapped resulting contigs to probe sequences and mined published genomes for outgroup and one ingroup sequence using the Phyluce pipeline (Faircloth, 2015), aligned orthologous sequences with Upp (Nguyen et al., 2015), and trimmed using Gblocks (Talavera and Castresana, 2007) with stringency settings relaxed from the default. I discarded any sequences with fewer than 114 taxa (70%) from further analyses. This resulted in 2,166 orthologous loci, 412 bases long on average.

Extraction of protein-coding sequences was done by blasting (Camacho et al., 2009) UCE loci against published proteins of three reference ant species (*Acromyrmex echinatior*, *Harpegnathos saltator*, and *Ooceraea biroi*), followed by collecting of protein queries and their nucleotide equivalents from best BLASTX hits for each locus using a custom bioinformatics pipeline. Separate alignment and trimming of resulting data set to include no fewer than 70% of all taxa produced an amino acid matrix of 1,103 loci 89.5 k amino acids long.

### Phylogenetics

I first estimated a gene tree for each locus using RAxML (Stamatakis, 2014) under GTR+4Γ model with 200 rapid bootstrap replicates. I then used individual locus characteristics to assess whether loci with different properties produce different phylogenies. Most basic statistics, such as alignment length, proportion of variable sites, and missing data were calculated using AMAS (Borowiec, 2016a). Using a custom R script I computed average branch lengths and average bootstrap support, used here as a proxy for rate of evolution and phylogenetic signal, respectively. With a custom Python script I calculated RCFV (Zhong et al., 2011), a measure of compositional heterogeneity, and performed a simulation-based compositional heterogeneity test with the Python p4 phylogenetics toolkit (Foster, 2004). Following this, I constructed a concatenated matrix of 271 loci with highest average bootstrap support, equal in length to 1/5 sites present in the combined data set. I also prepared a matrix of 379 compositionally homogeneous loci only and an alignment of identical to the combined data matrix but with *Aenictus* removed. I also constructed an amino acid matrix from extracted protein-coding sequences. For each of the concatenated nucleotide UCE data sets I used three partitioning schemes: 1) partitioning by UCE locus, 2) partitioning using the k-means algorithm (Frandsen et al., 2015) implemented in PartitionFinder2 (Lanfear et al., 2017), and 3) an unpartitioned model. The protein-coding sequences were analyzed as partitioned by locus. I estimated the phylogeny with RAxML under the partitioned GTR+4Γ model and 500 rapid bootstrap replicates (see Extended Methods) for all nucleotide matrices except for combined data partitioned by locus, which was analyzed with 100 bootstrap replicates. The amino acid matrix was analyzed under JTT+4Γ model and 100 bootstrap replicates. In addition to maximum likelihood analysis on all matrices, I also ran Bayesian analysis using ExaBayes (Aberer et al., 2014) on combined dataset. Consistent with the recent criticism of the k-means algorithm (Baca et al., 2017), this partitioning scheme appears to result in incorrect topology under ML analysis of the combined data and “high signal” loci matrices but not in analyses of slow-evolving or compositionally homogeneous loci or the Bayesian analysis of combined data. The concatenated analyses were done on CIPRES (Miller et al., 2010) and on the University of Rochester Center for Integrated Research Computing BlueHive computer cluster. I also used the statistical binning pipeline (Bayzid et al., 2015) and ASTRAL (Mirarab and Warnow, 2015) to estimate summary species trees on the combined data matrix, “high signal”, and slow-evolving loci.

### Divergence Time Estimation

I employed two different strategies to estimate the time-calibrated tree. The first approach used penalized likelihood and the fixed tree topology and branch length from the slow-evolving data. I used penalized likelihood as implemented in the chronos function of the R package ape (Paradis, 2013). I used an outgroup-rooted tree from the concatenated analysis of slow-evolving loci and included calibrations for nodes in the outgroup. Hard bounds were placed on node calibrations using fossil ages for the upper bound and previous estimates for node ages as lower bounds (For more details see Supplementary Table 3). The strict molecular clock could not be rejected for this tree, and I performed the analysis for 100 replicates in order to assess robustness to random starting values of the algorithm. The second approach utilized Bayesian inference with only ingroup taxa included, with several clades constrained as monophyletic and the root position and backbone relationships of the tree sampled from the posterior. Because Bayesian divergence time estimation is computationally very expensive, that approach was limited to 109 loci evenly distributed across the spectrum of rate of evolution. I used BEAST2 (Bouckaert et al., 2014) under the unpartitioned GTR+4Γ model, uncorrelated molecular clock with branch lengths drawn from a lognormal distribution, and fossilized birthdeath process (Heath et al., 2014) conditioning on the root to obtain divergence time estimates and a posterior sample of trees for biogeographic analyses. Calibrations included seven doryline fossils (see Supplementary Table 4 for more details): A *Neivamyrmex* sp. from Chiapas Amber, five species from Dominican Amber (*Acanthosticus hispaniolicus*, *Cylindromyrmex inopinatus*, *C. electrinus*, *C. antillanus*, and *Neivamyrmex ectopus*), and *Chrysapace* sp. from Baltic Amber. Another Baltic Amber fossil, *Procerapachys* spp. was not used because of uncertainty in its placement among crown or stem dorylines. I ran four independent MCMC chains for over 250 million generations until convergence was reached. Combined effective sample size (ESS) for all parameters was above 200.

### Diversification Analyses

To assess diversification rate shifts on the doryline phylogeny I used BAMM v2.5 (Rabosky, 2014). For input I used the consensus tree from BEAST analyses and sampling probabilities based on extant species diversity estimates (Borowiec, 2016b) for each genus in order to correct for uneven sampling of extant taxa across the phylogeny. I ran the MCMC for 20 million generations, sampling every 2,000 generations.

### Biogeographic History Estimation

I used the R package BioGeoBEARS (Matzke, 2013, 2014) for model selection and estimation of biogeographic history of the dorylines. As input I used the consensus tree from BEAST runs and a sample of 100 trees from the posterior of the same analysis. I compared six models commonly used for biogeographic inference on the consensus tree, with the DEC+J model emerging as the best-fitting. I then estimated biogeographic history on the 100 trees under this model to account for uncertainty in the deeper relationships and the timeline of early divergences. These results were then summarized on the consensus tree.

### Extended Methods

#### Data availability

Trimmed reads generated for this study are available at the NCBI Sequence Read Archive (to be submitted upon acceptance for publication). Sequence files, alignments, configuration files, and output of analyses, including phylogenetic trees, are available on Zenodo: https://doi.org/10.5281/zenodo.569071. Custom scripts used in this study are available on GitHub: https://doi.org/10.5281/zenodo.571246. Pipeline for extracting protein-coding sequences is available in a separate GitHub repository at https://doi.org/10.5281/zenodo.571247.

#### Taxon sampling

Taxon sampling included 154 newly sequenced ingroup species from all 27 currently recognized genera of Dorylinae ants and was guided by previous taxonomic and phylogenetic studies (De Andrade, 1998a; Borgmeier, 1955; Borowiec and Longino, 2011; Brady, 2003; Brady and Ward, 2005; Brady et al., 2014; Jaitrong and Yamane, 2011; MacKay, 1996; Bolton and Fisher, 2012) and expertise acquired preparing the global genus-level taxonomic revision of the group (Borowiec, 2016b). In addition to sampling all doryline genera, species from all biogeographic regions (as defined in (Cox, 2001) but treating the Malagasy region separate from Afrotropical) were included for each major lineage. Nine outgroup and one ingroup species (*Ooceraea biroi*) were also included based on publicly available ant genomes: *Atta cephalotes* (Suen et al., 2011), *Camponotus foridanus* (Bonasio et al., 2010), *Cardiocondyla obscurior* (Schrader et al., 2014), *Harpegnathos saltator* (Bonasio et al., 2010), *Linepithema humile* (Smith et al., 2011a), *Ooceraea biroi* (Oxley et al., 2014), *Pogonomyrmex barbatus* (Smith et al., 2011b), *Solenopsis invicta* (Wurm et al., 2011), and *Vollenhovia emeryi* (Smith et al., 2015).

#### Molecular data collection and sequencing

I extracted DNA from all newly sequenced specimens (Supplementary Table 1) using a DNeasy Blood and Tissue Kit (Qiagen, Valencia, CA, USA). Most specimens were extracted non-destructively, with extraction voucher retained. For several extractions the DNA collection was done destructively and a voucher specimen from the same colony was kept. I quantified DNA for each extraction using a Qubit fluorometer (Thermo Fisher Scientific, Waltham, MA, USA) and sheared *<*5–50 ng of DNA to a target size of approximately 400–600 bp. The shearing was done by sonication on a Bioruptor machine (Diagenode Inc., Philadelphia, PA, USA).

The library preparation protocol that follows was slightly modified from Blaimer et al. (2015). I used a KAPA Hyper Prep Library Kit (Kapa Biosystems, Inc., Wilmington, MA, USA) with magnetic bead cleanup (Fisher et al., 2011) and a SPRI substitute (Rohland and Reich, 2012) as described in (Faircloth et al., 2014). I used TruSeq adapters (Faircloth and Glenn, 2012) for ligation followed by PCR amplification of the library using a mix of HiFi HotStart polymerase reaction mix (Kapa Biosystems), Illumina TruSeq primers, and nuclease-free water. The following thermal cycler program was used for the PCR: 98 °C for 45 s; 13 or 14 cycles of 98 °C for 15 s, 65 °C for 30 s, 72 °C for 60 s, and final extension at 72 °C for 5 m. After rehydrating in 23 μl pH 8 Elution Buffer (EB hereafter) and purifying reactions using 1.1–1.2× speedbeads, I pooled nine to eleven libraries at equimolar ratios for final concentrations of 132–212 n/μl.

I enriched each pool with 9,446 custom-designed probes (MYcroarray, Inc.) targeting 2,524 UCE loci in Hymenoptera (Branstetter et al., 2017). I followed library enrichment procedures for the MYcroarray MYBaits kit (Blumenstiel et al., 2010), except I used a 0.1 *×* of the standard MYBaits concentration, and added 0.7 μl of 500 μmol custom blocking oligos designed against the custom sequence tags. I ran the hybridization reaction for 24 h at 65 °C, subsequently bound all pools to streptavidin beads (MyOne C1; Life Technologies), and washed bound libraries according to a standard target enrichment protocol (Blumenstiel et al., 2010). I used the with-bead approach for PCR recovery of enriched libraries as described in (Faircloth, 2015). I combined 15 μL of streptavidin bead-bound, enriched library with 25 μL HiFi HotStart Taq (Kapa Biosystems), 5 μL of Illumina TruSeq primer mix (5 μmol each) and 5 μL of nuclease-free water. Postenrichment PCR had the following profile: 98 °C for 45 s; 18 cycles of 98 °C for 15 s, 60 °C for 30 s, 72 °C for 60 s; and a final extension of 72 °C for 5 m. I purified resulting reactions using 1.1–1.2 × speedbeads, and rehydrated the enriched pools in 22 μL EB.

Following enrichment I quantified 2 μL of each pool using a Qubit fluorometer (broad range kit). I verified if the enrichment was successful by amplifying four UCE loci targeted by the probe set. I set up a relative qPCR by amplifying two replicates of 1 ng of enriched DNA from each pool for the four loci and comparing those results to two replicates of 1 ng unenriched DNA from each pool. I performed qPCR using a SYBR FAST qPCR kit (Kapa Biosystems) on CFX Connect Real-Time PCR Detection System (Bio-Rad). Following data collection, I calculated fold-enrichment values, assuming an efficiency of 1.78 and using the formula 1.78 × abs(*enrichedCp* – *unenrichedCp*). I then performed qPCR quantification by creating dilutions of each pool (1:200,000, 1:800,000, 1:1.000,000, 1:10.000,000) and assuming an average library fragment length of 600 bp. Based on the size-adjusted concentrations estimated by qPCR, I pooled libraries at equimolar concentrations.

The pooled libraries were then subjected to further quality control on Bioanalyzer and sequenced using one full and one partial lane of a HiSeq 125 Cycle Paired-End Sequencing v4 run. QC and sequencing were performed at the University of Utah High Throughput Genomics Core Facility. Quality-trimmed sequence reads generated as part of this study are available from the NCBI Sequence Read Archive (to be submitted upon acceptance for publication).

#### Processing of UCE data

Read cleaning, assembly, matching of contigs to probes, construction of the unaligned data set, and alignment trimming were done using the Phyluce pipeline scripts (Faircloth, 2015). I trimmed the FASTQ data using `illumiprocessor`, a wrapper around Trimmomatic (Bolger et al., 2014), with default settings (`LEADING:5, TRAILING:15, SLIDINGWINDOW:4:15, MINLEN:40`). Assemblies were done using Trinity v20140717 (Grabherr et al., 2011) with the `phyluce_assembly_assemblo_trinity` wrapper. The orthology assessment was then done by matching the assembled contigs to enrichment probe sequences with `phyluce_assembly_match_contigs_to_probes` (min_coverage=50, min_identity=80). This step generated a `sqlite` database which was then used to build FASTA files for the 2,524 orthologous loci with `phyluce_assembly_get_match_counts`, `phyluce_assembly_get_fastas_from_match_counts`, and `phyluce_assembly_explode_get_fastas_file`.

#### Extraction of protein-coding sequences

For the purpose of extracting protein-coding data from the sequenced UCE loci I developed a custom bioinformatics workflow that consists of three major components: using NCBI BLASTX (Camacho et al., 2009) to match unaligned UCE sequences to reference proteins and 2) choosing the best hit for each sequence followed by 3) extraction of protein queries and their nucleotide equivalents from those hits. Using `makeblastdb` program of the BLAST package I prepared a database from three publicly available collections of protein sequences of *Acromyrmex echinatior* (Nygaard et al., 2011), *Harpegnathos saltator* (Bonasio et al., 2010), and *Ooceraea biroi* (Oxley et al., 2014). Using BLASTX against this database resulted in one BLAST output file per UCE locus, containing multiple matches (hits) for each UCE sequence (taxon). Each hit, in turn, may be composed of one or more ranges that correspond to protein fragments (exons). I used BLAST scores for those ranges to identify best hits for each taxon and UCE. This was done with custom Python code using Biopython’s (Cock et al., 2009) module for parsing BLAST XML output and the following logic:

For each sequence (taxon) and hit within, both total and maximum scores are tallied. If a hit’s total score is equal to the maximum score of its ranges and it corresponds to the maximum score for the taxon, such hit is considered best and is kept. This means that this hit was composed of a single range and its score was not exceeded by any other hits, composed of one or multiple ranges. If a hit’s total score is higher than the maximum score of any one of its ranges, and that hit’s total score is the best hit score for a species, this hit is kept unless it contains overlapping ranges. Finally, if the total hit score is equal to its maximum score but not to the best hit score for a species, its total score is checked to see if it corresponds to the highest individual range score for a species. If this is true, the hit is kept. Such hit would be composed of single range and considered best even if hits with higher total scores but overlapping ranges are present. If composed of multiple ranges, a best hit is concatenated into a query in the order based on its coordinates and its presence on either forward or reverse strand. These protein queries are then matched to corresponding input nucleotide sequence or its reverse complement.

If introns that do not change reading frame are present, translations of the query sequence may span across them. Because of this I performed additional trimming if long (4 sites or more) gaps in the subject protein sequence were found. All sites corresponding to those long gaps were trimmed from the protein query and its nucleotide equivalent. If at this point there is still a stop codon in the protein query, such record was discarded.

For each record the resulting protein queries and their corresponding nucleotides were considered ready for downstream analyses.

#### Alignment and trimming

Assembly and contig matching resulted in sequences of varying lengths across taxa, as evidenced by summaries produced with `phyluce_assembly_get_fasta_lengths`. Because of this I used UPP (Nguyen et al., 2015), a phylogeny-aware alignment tool designed to align fragmentary sequences to a backbone of longer sequences. Based on the performance (recovered final post-trimming alignment length) of different settings, for the backbone I chose the cutoff of 30% of the longest sequences present in each locus. Because the UPP version used did not have an option to specify a fixed number of longest sequences in the backbone, a locus-specific command was printed for each locus based on its fragment size distribution with UPP’s `-M` option set to longest sequence and threshold (option `-T`) calculated to encompass 30% of taxa. UPP was also set to filter sequences from the backbone if their branches were 5 times longer than the median for all backbone sequences (`-l 5` argument):

~~~
run_upp.py -s [input_alignment] -M [longest_sequence_length]
-T [locus_threshold] -d [alignment_output_directory] -o [output]
-p [temporary_output_directory] -l 5
~~~

These custom commands were printed with `print_upp_command.py`, a custom script utilizing code from `phyluce_assembly_get_fasta_lengths`.

Although alignment trimming has been recently criticized (Tan et al., 2015), the untrimmed alignments contained on average more than 75% of gaps and missing data. Because of the substantial computational burden of gap-rich data analysis, I trimmed the alignments using Gblocks (Talavera and Castresana, 2007) under settings relaxed from the default (`b1=0.5 b2=0.5 b3=12 b4=7`). I calculated alignment statistics and manipulated the files using AMAS v0.98 (Borowiec, 2016a). All loci with fewer than 114 taxa (less than 70%) were discarded, resulting in 2,166 out of 2,524 loci for downstream analyses. These loci had on average 151.7 (92.5%) taxa, were 412.2 nucleotides long, and had 7.7% missing data (gaps). Due to computing time constraints, loci with protein-coding sequences extracted were aligned using MAFFT v7.300b with `--leavegappyout` setting turned on. These alignments were trimmed using Gblocks with settings as above and further trimmed for outlier taxa using a custom R script and AMAS, removing any ingroup sequences whose uncorrected p-distance was more than 3σ from the mean for a locus.

#### Partitioning

PartitionFinder 2 (Lanfear et al., 2017) was used to partition concatenated alignments using the k-means clustering of sites based on evolutionary rates (Frandsen et al., 2015). A starting tree for model fit and site clustering algorithm was generated with RAxML Pthreads v8.2.3 (Stamatakis, 2014) using the fast tree search algorithm (`-f E` flag). Because of recent criticism of the k-means algirithm (Baca et al., 2017), I performed additional ML analyses for combined data set under unpartitioned and partitioned by locus schemes. Protein-coding sequences were analyzed as partitioned by locus. K-means tends to result in relatively low number of partitions and other partitioning by locus scheme was not computationally feasible for Bayesian inference.

#### Phylogenetic analyses using maximum likelihood

For each locus I estimated a gene tree with RAxML Pthreads v8.2.3 under a general time-reversible model of sequence evolution with rate modeled using a gamma distribution discretized into four bins (GTR+4Γ model). 200 rapid bootstraps were followed by a thorough search of the maximum likelihood tree:

~~~
raxml -T [no_cores] -m GTRGAMMA -f a -# 200 -p 12345 -x 12345
-s [input_alignment] -n [output_file_name]
The same mode of inference was applied to supergenes created by the statistical binning pipeline (see Species Tree analyses below) but each supergene was partitioned by constituent loci using a partitions file (-q) and the -M flag was added for a fully partitioned analysis with branch lengths optimized separately for each partition (Bayzid et al., 2015):
raxml -T [no_cores] -m GTRGAMMA -f a -# 200 -p 12345 -x 12345 -M
-s [input_alignment] -q [partitions_file] -n [output_file_name]
~~~

I ran the analyses of concatenated matrices with RAxML Hybrid v8.2.4 and v8.2.9 on CIPRES Portal (Miller et al., 2010) using the same model and bootstrap settings but with a pre-defined partitioning scheme (see above), no `-M` option due to the high number of partitions, and either 100 (amino acid analysis and combined data partitioned by locus) or 500 bootstrap replicates (all other analyses). The amino acid analysis was ran under the JTT+4Γ model.

#### Phylogenetic analyses using Bayesian inference

I used ExaBayes v1.4.1 (Aberer et al., 2014) to perform analysis on k-means partitioned matrix of combined data set under GTR+4Γ. The analysis was ran with two runs, four MCMC chains each for 5 million generations. I disabled the default of parsimony starting tree such that analysis was initiated with a random topology. Convergence and mixing of the MCMC were determined by monitoring average standard deviation of split frequencies, which are considered acceptable below 5% (final value 1.75% for combined data set) and effective sample sizes (ESS) for all parameters, considered acceptable above 200 (min ESS: 900). I used 25% burn-in to construct consensus trees.

#### Species tree analyses

In addition to phylogenetic inference on concatenated loci I performed species tree analyses that attempt to reconcile gene tree incongruencies arising due to incomplete lineage sorting (Edwards, 2009). I used Accurate Species TRee Algorithm, ASTRAL v4.10.12 (Mirarab and Warnow, 2015; Sayyari and Mirarab, 2016). I used a weighted statistical binning pipeline (Bayzid et al., 2015) to create supergene alignments and trees. Locus trees used as input for the pipeline were considered under 75 bootstrap support threshold. Summary methods for species tree inference such as those used here have been shown to be negatively impacted by error in estimated input gene trees (Roch and Warnow, 2015). The binning approach was devised to alleviate this (Mirarab et al., 2014). The data sets analyzed with species tree approaches included binned supergenes of all loci (514 supergenes), supergenes identified in the high bootstrap loci (147 supergenes), and supergenes from the slow-evolving loci (100 supergenes).

#### Measures of compositional heterogeneity

Most models commonly used for phylogenetic inference, including the partitioned GTR+4Γ model used here, assume that alignments are compositionally homogeneous among taxa (Moran et al., 2015). To quantify among-taxon compositional heterogeneity of the data, I used two approaches: 1) statistical tests of compositional heterogeneity and 2) a continuous measure of relative composition frequency variability (RCFV) (Zhong et al., 2011). The former included a phylogeny-corrected statistical test that compares compositions in data sets simulated under a model (the null distribution) to the compositions in the observed alignment (Foster, 2004). For the purpose of this study the test was done on 200 simulated alignments for each observed alignment, assuming a GTR+4Γ model and a neighbor joining tree calculated using BioNJ (Gascuel, 1997). The often used but less appropriate *χ*^2^ test for compositional heterogeneity was also performed for comparison. The two tests were carried out using the p4 program for phylogenetic inference (Foster, 2004). Relative composition of frequency variability (RCFV) is the other measure used here to compare compositional heterogeneity among data (Zhong et al., 2011). RCFV is the sum of absolute values of differences observed among frequencies of all four nucleotides, divided by the number of taxa. The differences are calculated by subtracting the overall frequency of the character in a matrix from the frequency of that character in an individual sequence (taxon). The sum of these differences is then divided by the number of taxa and this number in turn is summed for each sequence/taxon in the alignment:

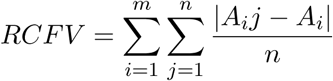

where *m* is the number of distinct character states (four for nucleotide sequences), *A*_*i*_*j* is the frequency of nucleotide *i* in taxon *j* and *A*_*i*_ is the frequency of character (nucleotide) *i* across *n* taxa. RCFV thus gives a relative measure of compositional heterogeneity for a data set, and as the sum of differences between frequencies is calculated for each sequence, it also allows for comparison among taxa within an alignment.

#### Tree-based locus statistics

Following maximum likelihood estimation of gene trees, I computed average branch length for each of the 2,166 loci. The average branch length is a proxy for the rate of evolution of each locus. Saturation is another property that is potentially correlated with poor model fit. This I calculated by plotting p-distances of an alignment against distances on the tree from model-based maximum likelihood inference (Philippe and Forterre, 1999). These distances would show a perfect fit to simple linear regression in the absence of saturation. When there is a need of correction for multiple substitutions, however, the curve will depart from linearity. I sorted each locus by slope of regression for a relative measure of saturation.

I computed all the tree-based measures with a custom R (v3.0.2 (2013-09-25)) script leveraging packages `ape` v3.1-1 (Paradis et al., 2004) and `seqinr` v3.1-2 (Charif and Lobry, 2007), modified from code originally developed for Borowiec et al. (2015).

#### Divergence time estimation

To build a time-calibrated chronogram of the Dorylinae I used the R (v3.2.3 (Team, 2014) package `ape` v3.4 and its function `chronos` (Paradis et al., 2004; Paradis, 2013). `Chronos` uses the penalized likelihood method (Sanderson, 2002) and allows selection of the molecular clock model best fitting the data using an information criterion introduced in (Paradis, 2013). I used the maximum likelihood tree with branch lengths, rooted with *Harpegnathos saltator* estimated from slow-evolving loci partitioned under k-means as input. The method requires that nodes are calibrated with hard bounds of minimum and maximum ages. The calibration scheme is given in Supplementary Table 3. The information criterion implemented in `chronos` identified the strict molecular clock as the best fitting. Because unknown dates are initialized with a random algorithm it is possible to assess the robustness of the node ages to these initial ages by running multiple independent analyses. I ran 100 `chronos` replicates and summarized output trees using the `sumtrees.py` script distributed as a part of `Denropy` package v4.0.3, (Sukumaran and Holder, 2010). The summarized tree has mean branch lengths mean node ages, as well as uncertainty captured as node age ranges obtained across the 100 replicates. I visualized the tree with mean node ages and age ranges using FigTree v1.4.2. Because the information criterion implemented was shown to be poor at distinguishing the strict clock from a model with a small fixed number of rates (Paradis, 2013), I repeated the procedure for a discrete clock model with 10 categories. The results are presented in Supplementary Figure 19.

Because calibrations requiring hard minimum and maximum ages may be considered prone to bias and because penalized likelihood does not utilize sequence data, I also performed Bayesian divergence dating using the recently developed birth-death process (Heath et al., 2014). This method assumes no prior belief on calibrated node ages, instead relying on a single recovery age of a fossil that is assumed to be a descendant of the calibrated node. The method simulates tree topology via a birth-death process and treats fossils as a part of the diversification process with variable attachment points on the tree and fixed recovery ages. In addition to assuming no prior beliefs on calibrated node ages, this method is not limited to using only the oldest fossils known for a given node. For these analyses I used BEAST v2.3.2 (Bouckaert et al., 2014) with Sampled Ancestors package (Gavryushkina et al., 2014). Because Bayesian divergence time estimation is computationally expensive with 150+ taxa, these analyses were limited to 109 loci (5% of all loci) sampled at even intervals throughout the rate spectrum. I set up the BEAST analyses under unpartitioned GTR+4Γ model of sequence evolution and uncorrelated clock sampling rates from a lognormal distribution. The analysis was set up with four independent runs for >300 million generations. I determined convergence and adequate sample size (all parameters sampled at ESS > 200) using Tracer v1.5. The calibration scheme included seven fossils with fixed sampling times obtained by drawing a random number from a uniform distribution (`runif` function in R base) bounded by minimum and maximum ages of the deposit in which the fossil is found (deposit ages follow the Fossilworks website (Alroy, 2016); Supplementary Table 4). I used conditioning on the root age with a prior.

#### Diversification analyses

BAMM v2.5 (Rabosky, 2014) analyses used the consensus BEAST chronogram and a table of sampling probabilities based on extant species diversity estimates for each genus (Borowiec, 2016b). The sampling proportions were set as follows: *Acanthostichus*: 0.17, *Aenictogiton*: 0.2, *Aenictus*: 0.09, *Cerapachys*: 0.5, *Cheliomyrmex*: 0.6, *Chrysapace*: 0.8, *Cylindromyrmex*: 0.4, *Dorylus*: 0.2, *Eburopone*: 0.12, *Eciton*: 0.58, *Eusphinctus*: 0.5, *Labidus*: 0.6, *Leptanilloides*: 0.17, *Lioponera*: 0.08, *Lividopone*: 0.13, *Neivamyrmex*: 0.11, *Neocerapachys*: 0.4, *Nomamyrmex*: 1, *Ooceraea*: 0.2, *Parasyscia*: 0.08, *Simopone*: 0.18, *Sphinctomyrmex*: 0.4, *Syscia*: 0.13, *Tanipone*: 0.4, *Vicinopone*: 1, *Yunodorylus*: 0.4, *Zasphinctus*: 0.15. I used the “setBAMMpriors” function in the R package BAMMtools (Rabosky et al., 2014) to create priors used for the analysis. I ran the MCMC for 20 million generations, sampling every 2,000 generations. I checked the convergence and plotted the analysis results using BAMMtools and CODA (Plummer et al., 2006).

### Biogeographic analyses

I used the maximum likelihood functions available in BioGeoBEARS R package (Matzke, 2013, 2014) to compare fit and select from among models commonly used for estimation of biogeographic histories. I discretized species distributions into six biogeographic regions following (Cox, 2001) and treating Malagasy, as the seventh separate region. The regions included Neotropical, Nearctic, Palearctic, Afrotropical, Malagasy, Indomalayan, and Australasian. The boundary between Indomalayan and Australasian regions is the Wallace Line. I set up a time-stratified analysis that assumed different region adjacency and dispersal probabilities between 110-50 Ma and 50 Ma-present (M3_stratified-type model of Matzke (2014)) and compared likelihoods and AICc scores under six models: DEC, DEC+J, DIVALIKE, DIVALIKE+J, BAYAREALIKE, and BAYAREALIKE+J.

---

**Supplementary Figure 1:**
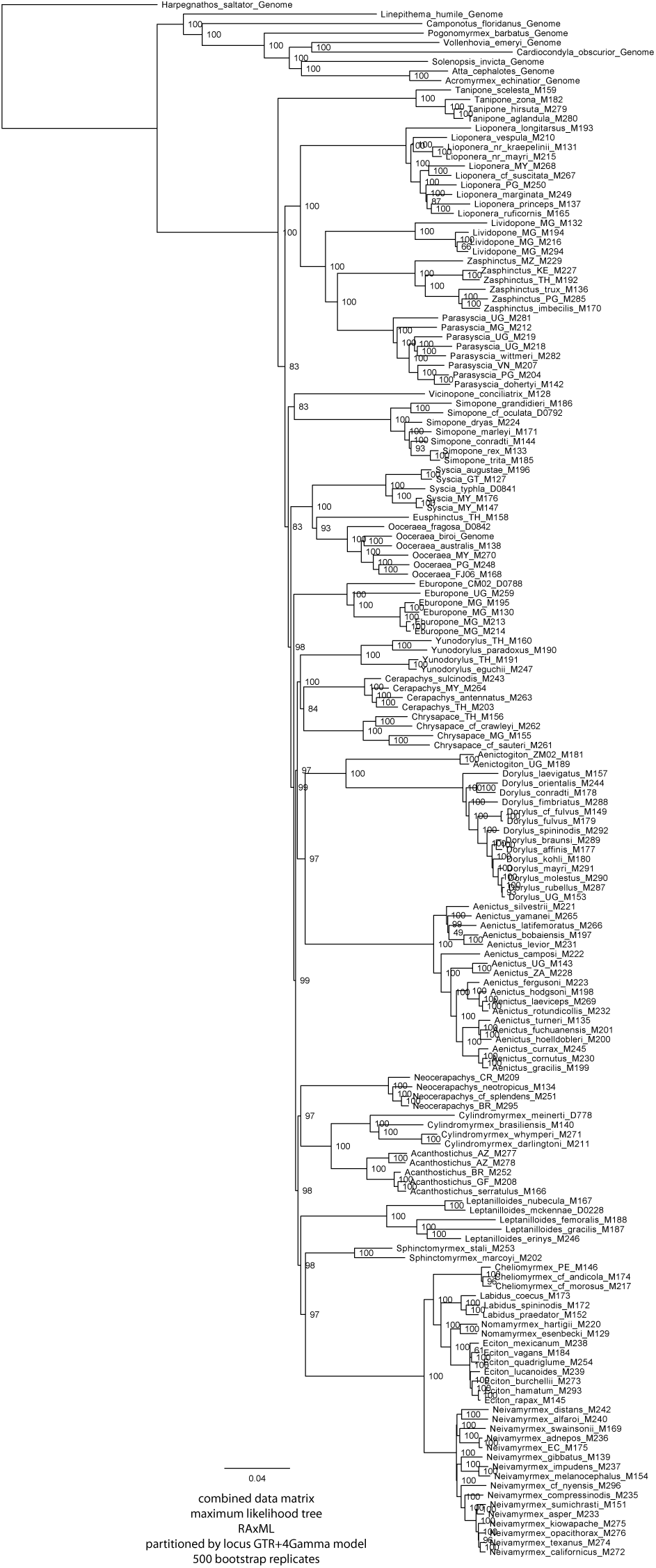
Maximum likelihood tree obtained from the combined data matrix partitioned by locus. Scale is in number of substitutions per site. Nodal support in percent bootstrap.

**Supplementary Figure 2:**
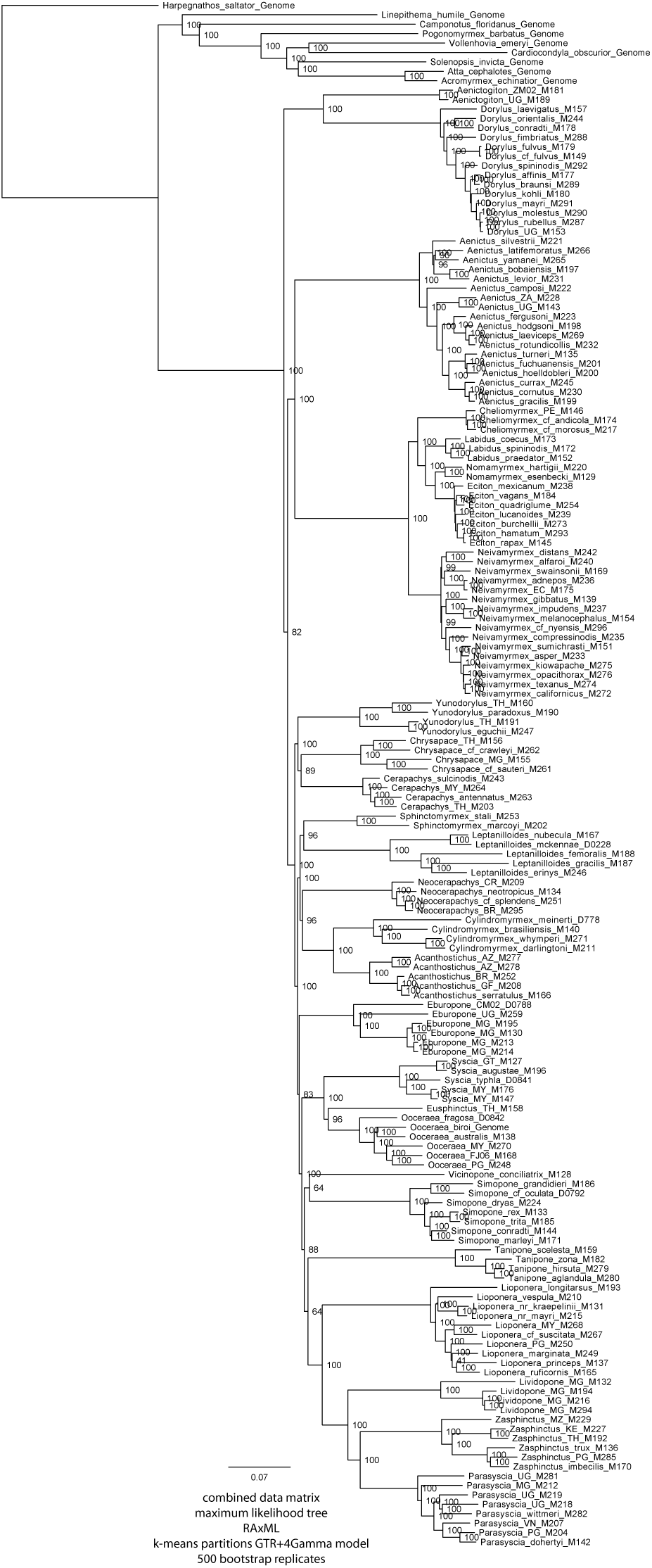
Maximum likelihood tree obtained from the combined data matrix under k-means partitions. Scale is in number of substitutions per site. Nodal support in percent bootstrap.

**Supplementary Figure 3:**
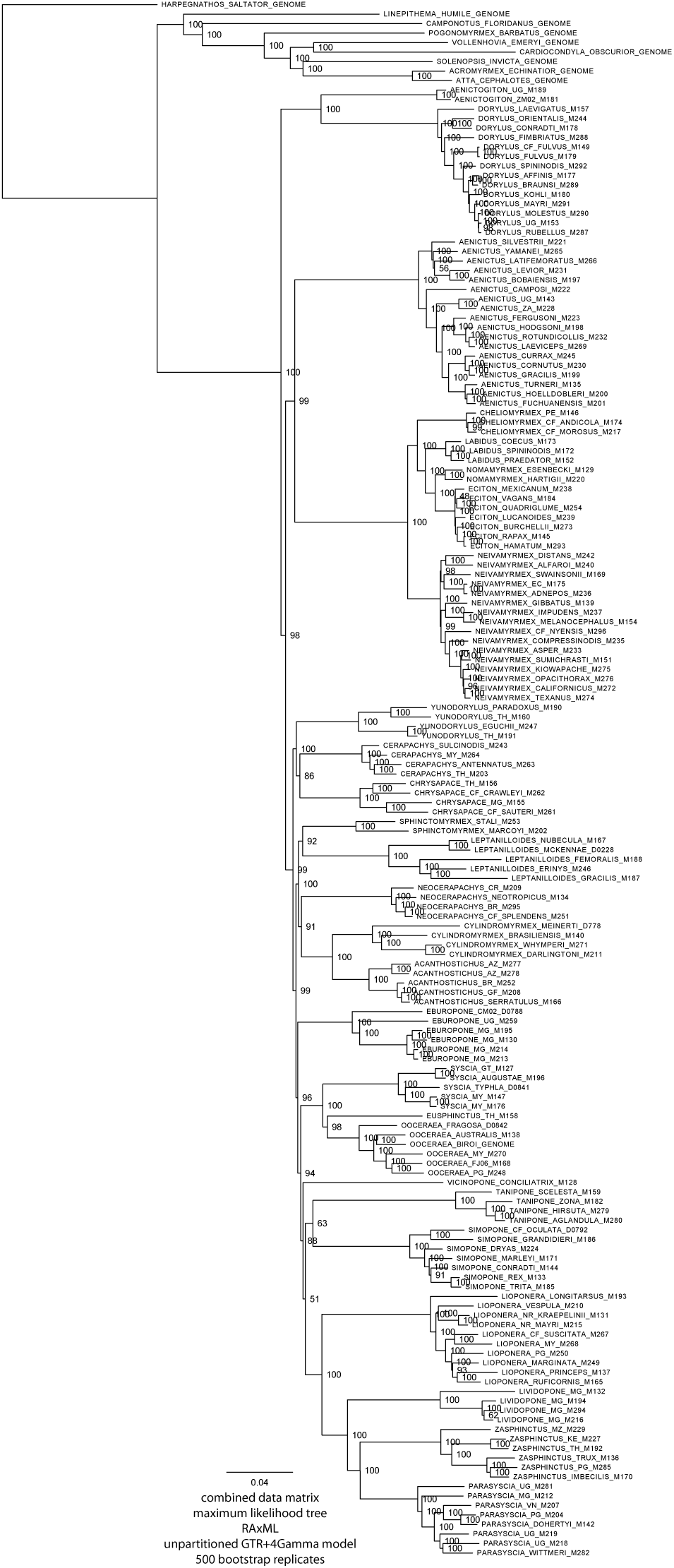
Maximum likelihood tree obtained from unpartitioned combined data matrix. Scale is in number of substitutions per site. Nodal support in percent bootstrap.

**Supplementary Figure 4:**
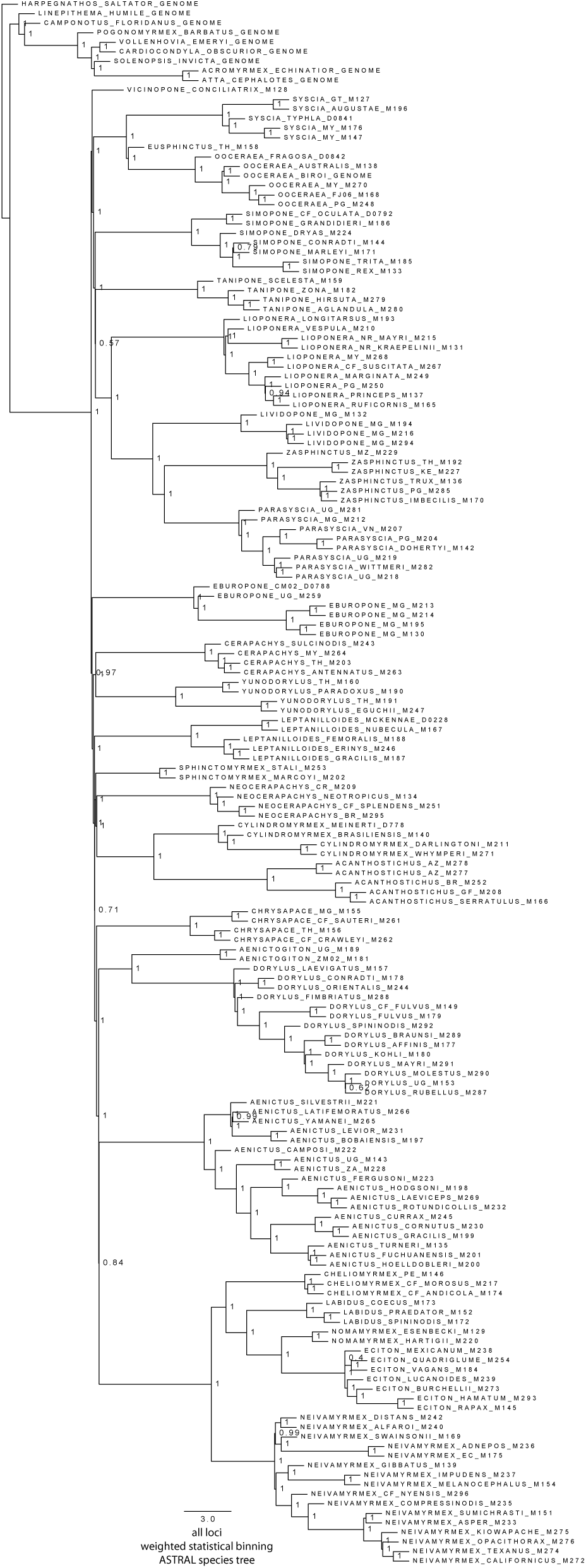
Summary species tree obtained from all 2,166 loci using ASTRAL. Scale is in coalescent units. Nodal support in local posterior probabilities.

**Supplementary Figure 5:**
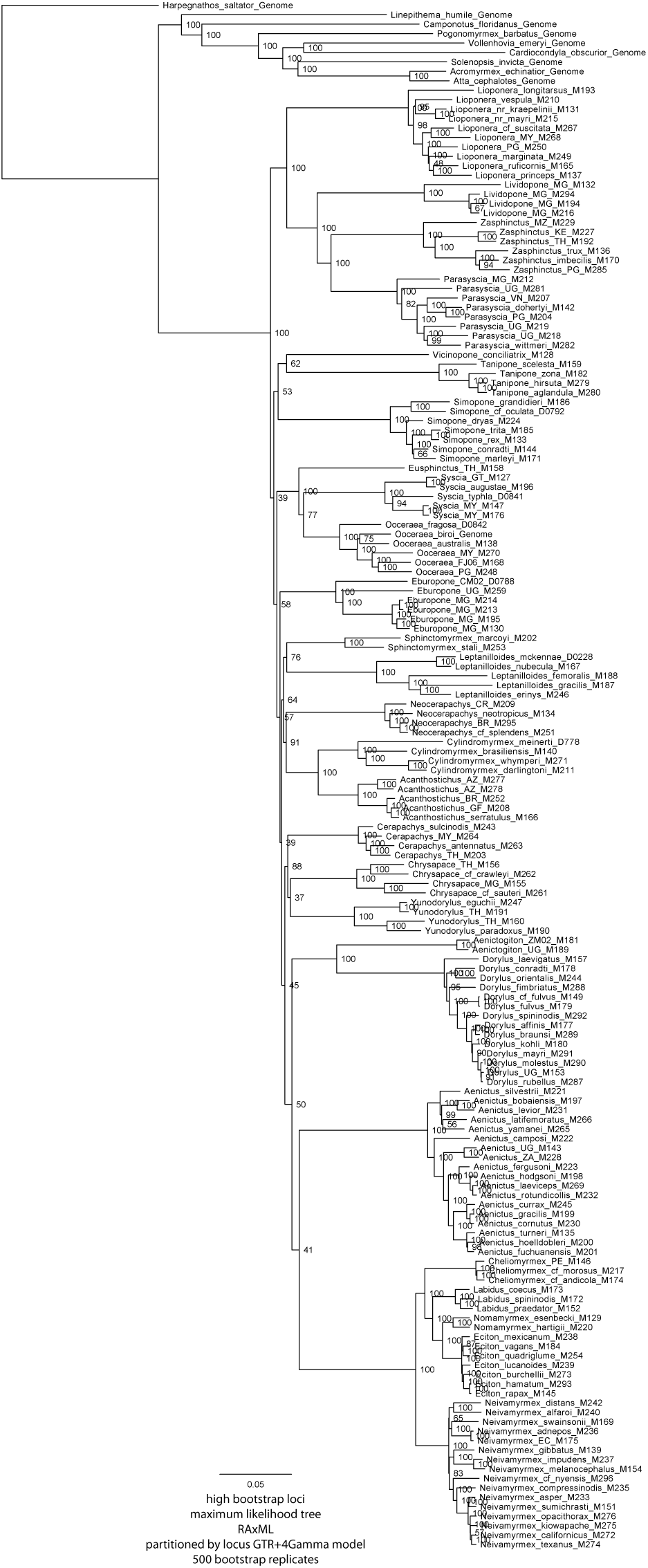
Maximum likelihood tree obtained from high-bootstrap data matrix partitioned by locus. Scale is in number of substitutions per site. Nodal support in percent bootstrap.

**Supplementary Figure 6:**
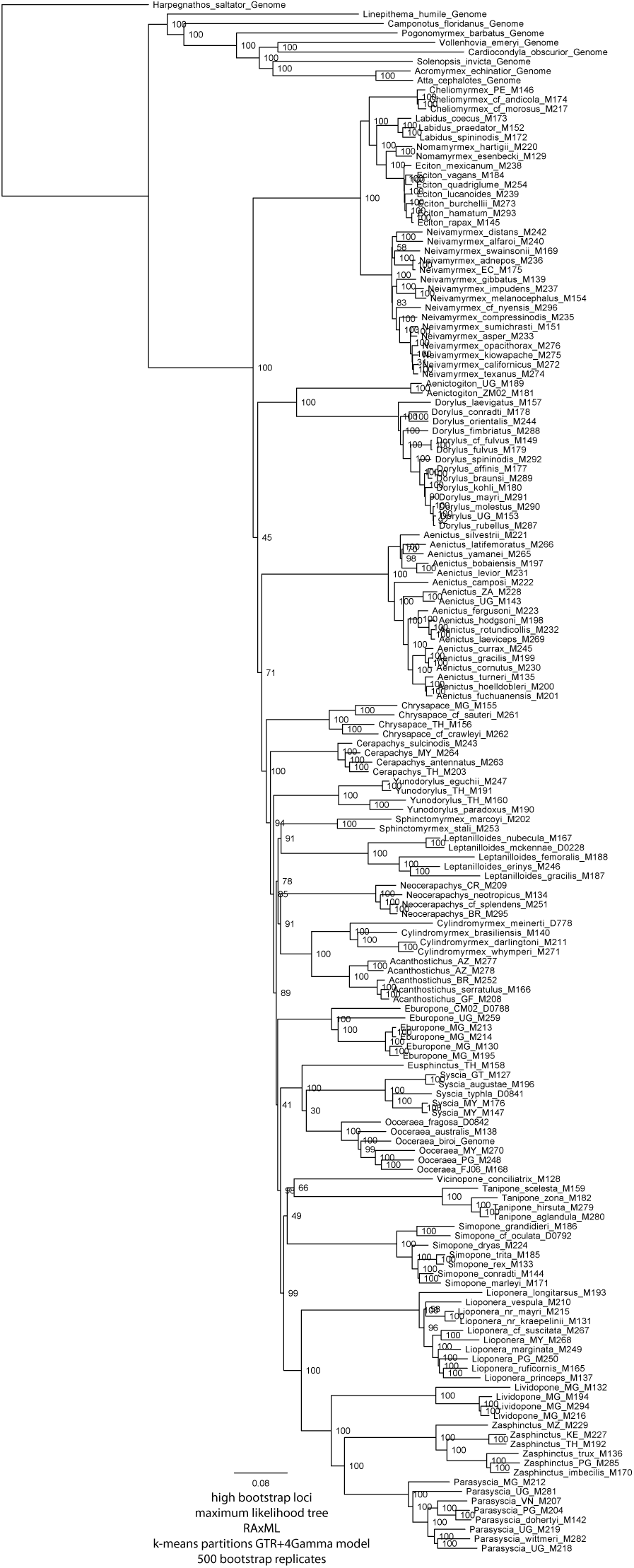
Maximum likelihood tree obtained from high-bootstrap data matrix under k-means partitions. Scale is in number of substitutions per site. Nodal support in percent bootstrap.

**Supplementary Figure 7:**
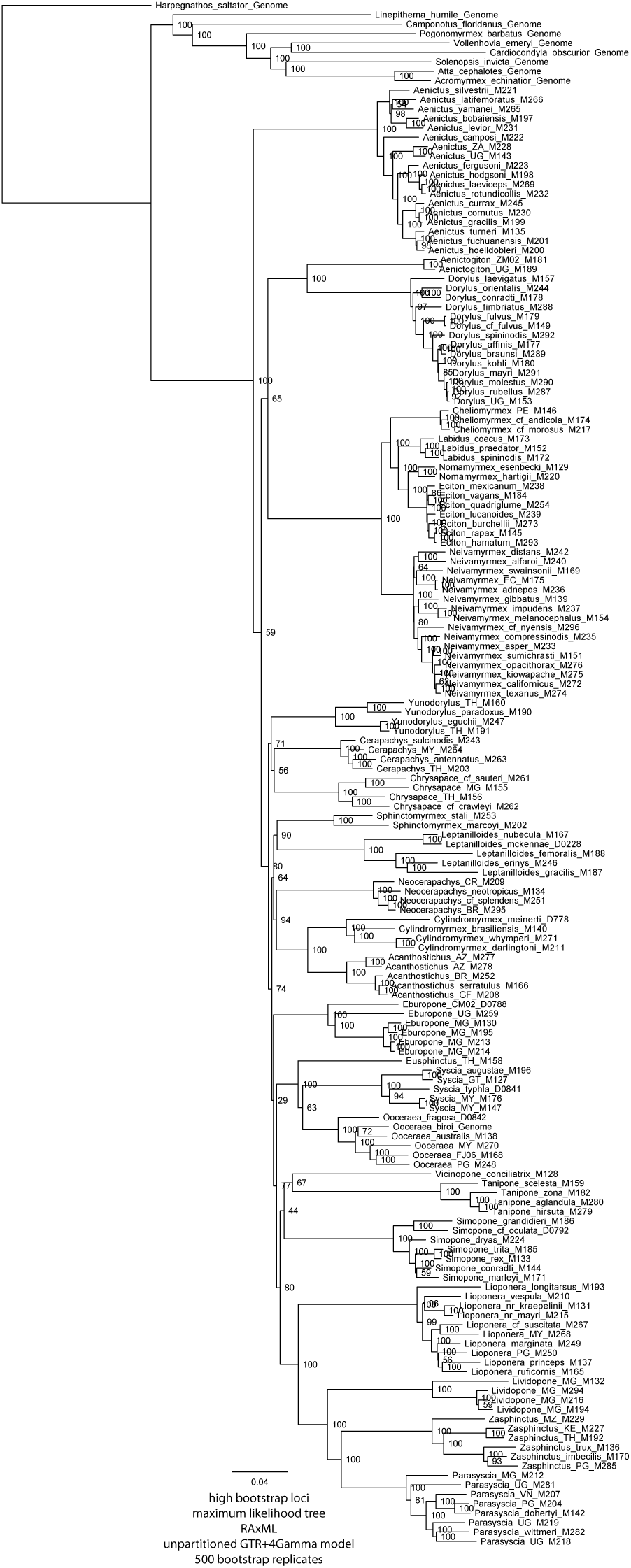
Maximum likelihood tree obtained from unpartitioned high-bootstrap data matrix. Scale is in number of substitutions per site. Nodal support in percent bootstrap.

**Supplementary Figure 8:**
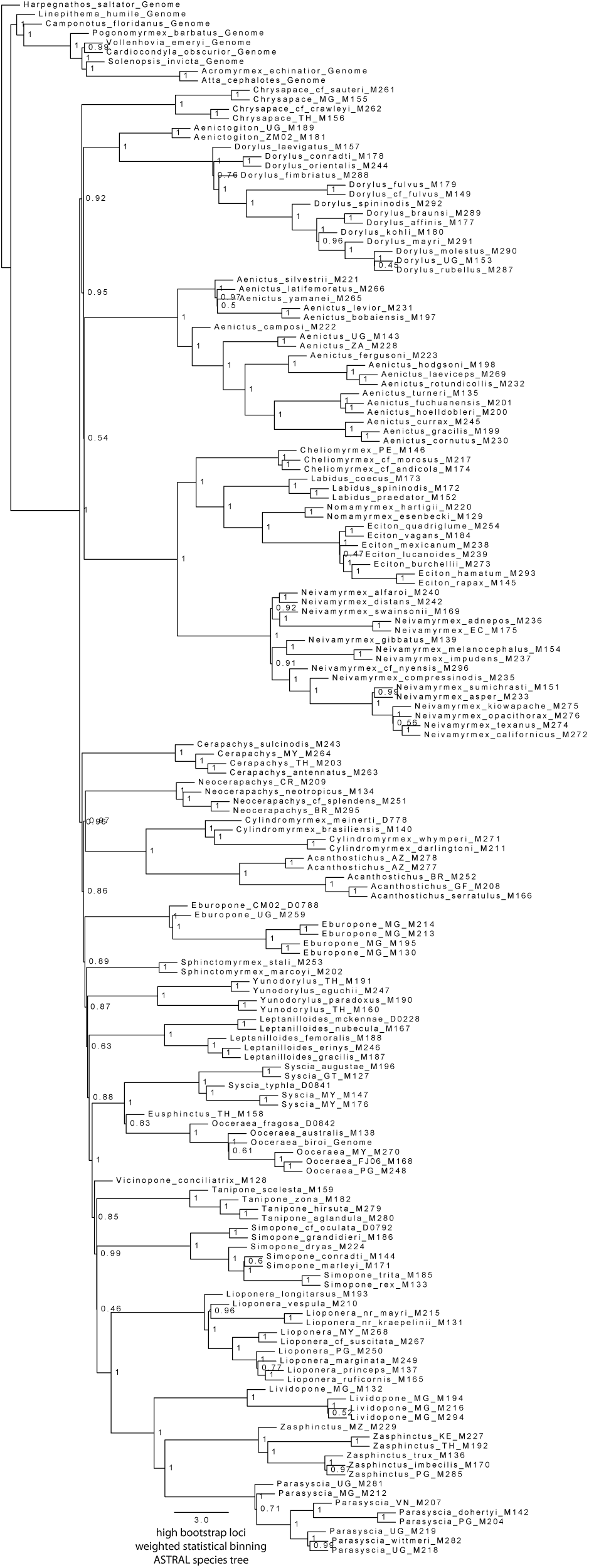
Summary species tree obtained from 271 highest bootstrap loci using ASTRAL. Scale is in coalescent units. Nodal support in local posterior probabilities.

**Supplementary Figure 9:**
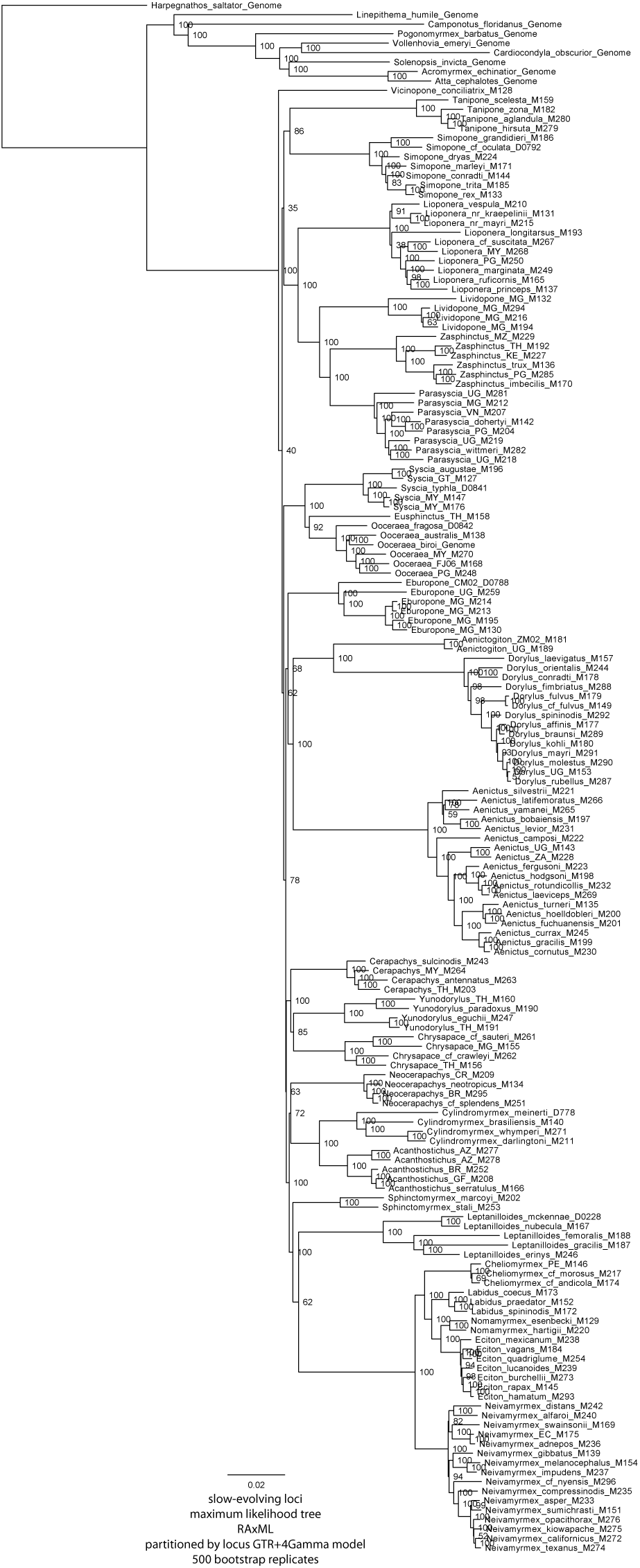
Maximum likelihood tree obtained from slow-evolving data matrix partitioned by locus. Scale is in number of substitutions per site. Nodal support in percent bootstrap.

**Supplementary Figure 10:**
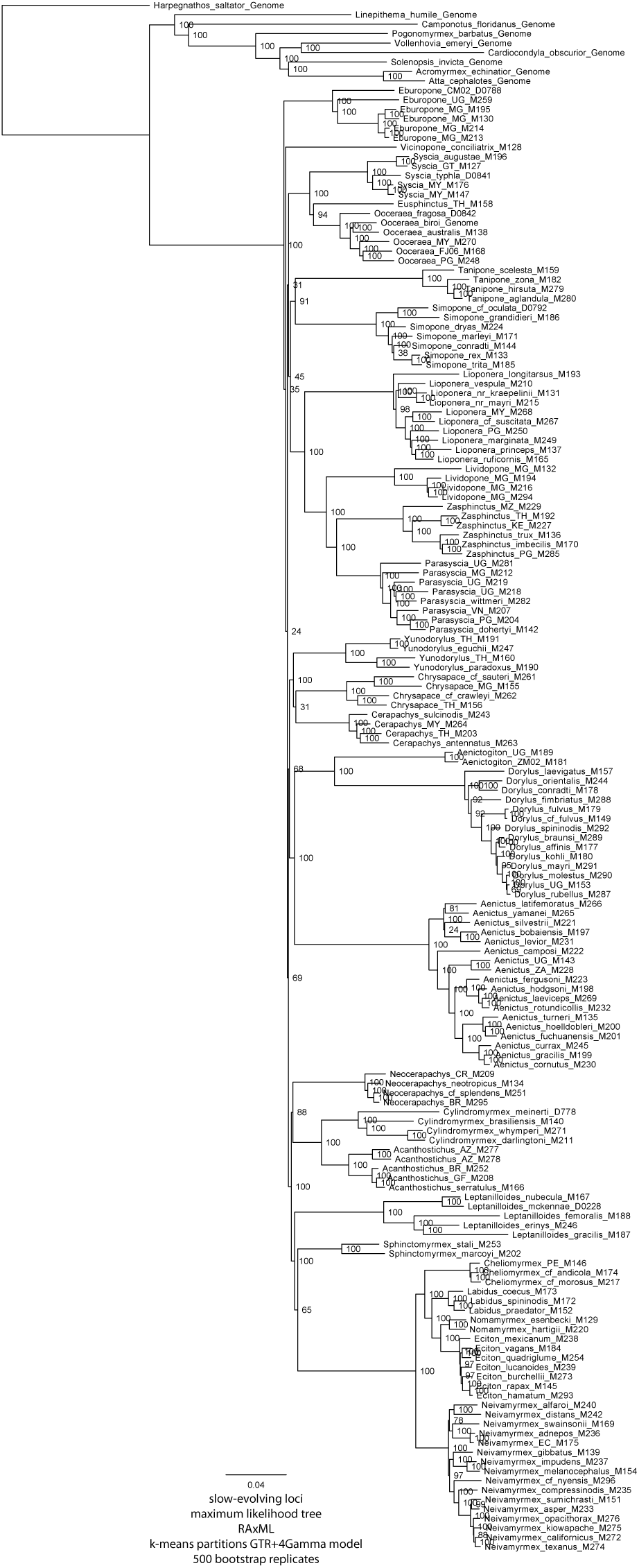
Maximum likelihood tree obtained from slow-evolving data matrix under k-means partitions. Scale is in number of substitutions per site. Nodal support in percent bootstrap.

**Supplementary Figure 11:**
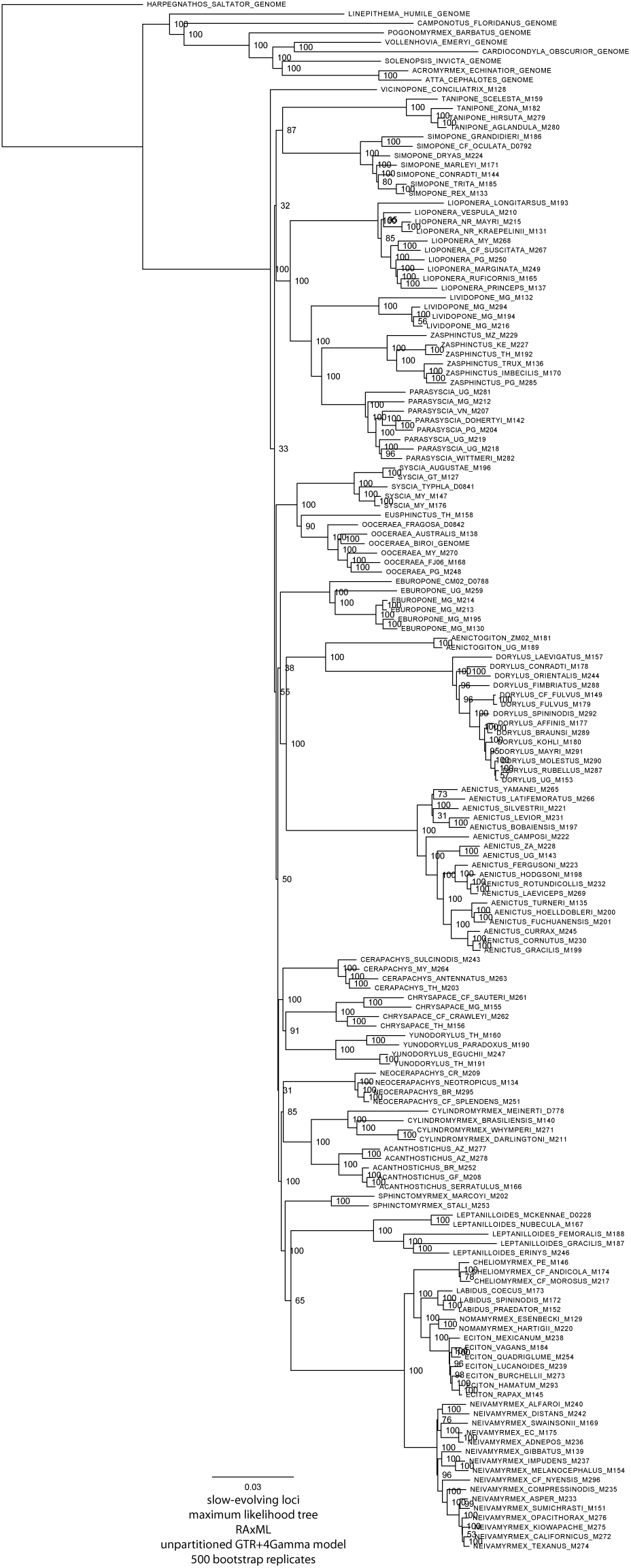
Maximum likelihood tree obtained from unpartitioned slow-evolving data matrix. Scale is in number of substitutions per site. Nodal support in percent bootstrap.

**Supplementary Figure 12:**
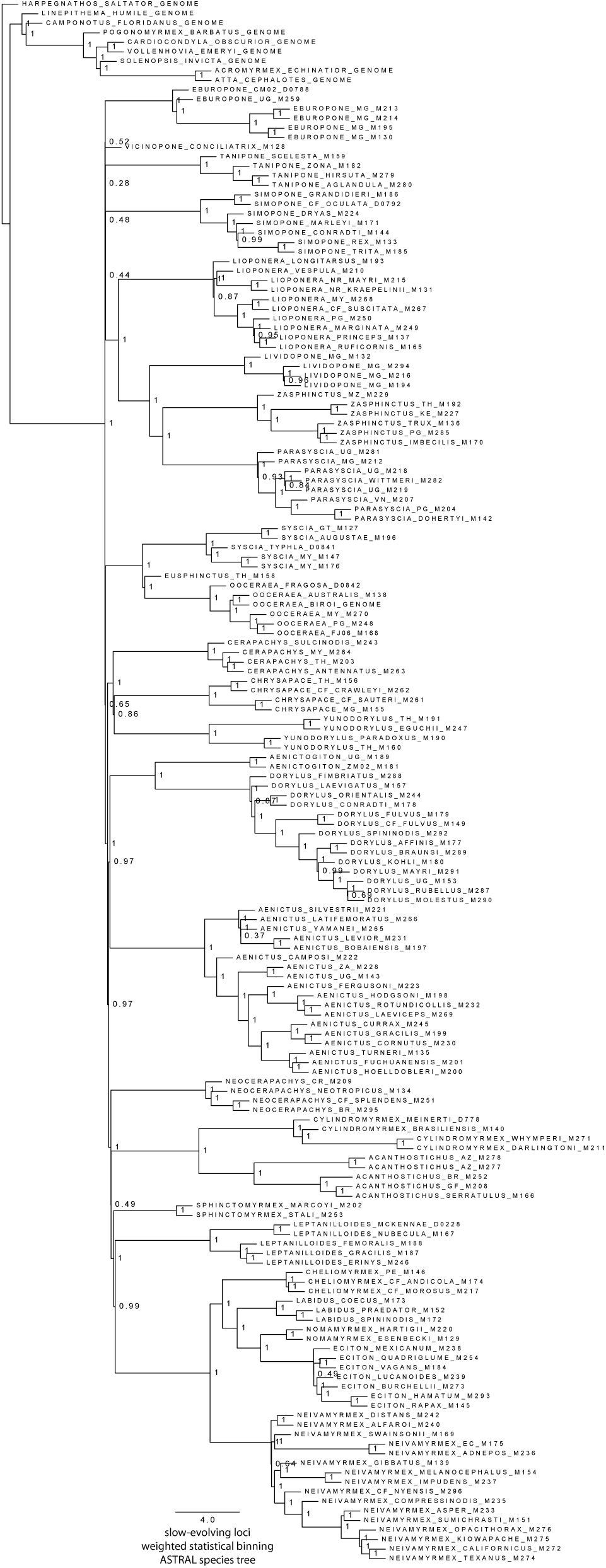
Summary species tree obtained from 580 slow-evolving loci using ASTRAL. Scale is in coalescent units. Nodal support in local posterior probabilities.

**Supplementary Figure 13:**
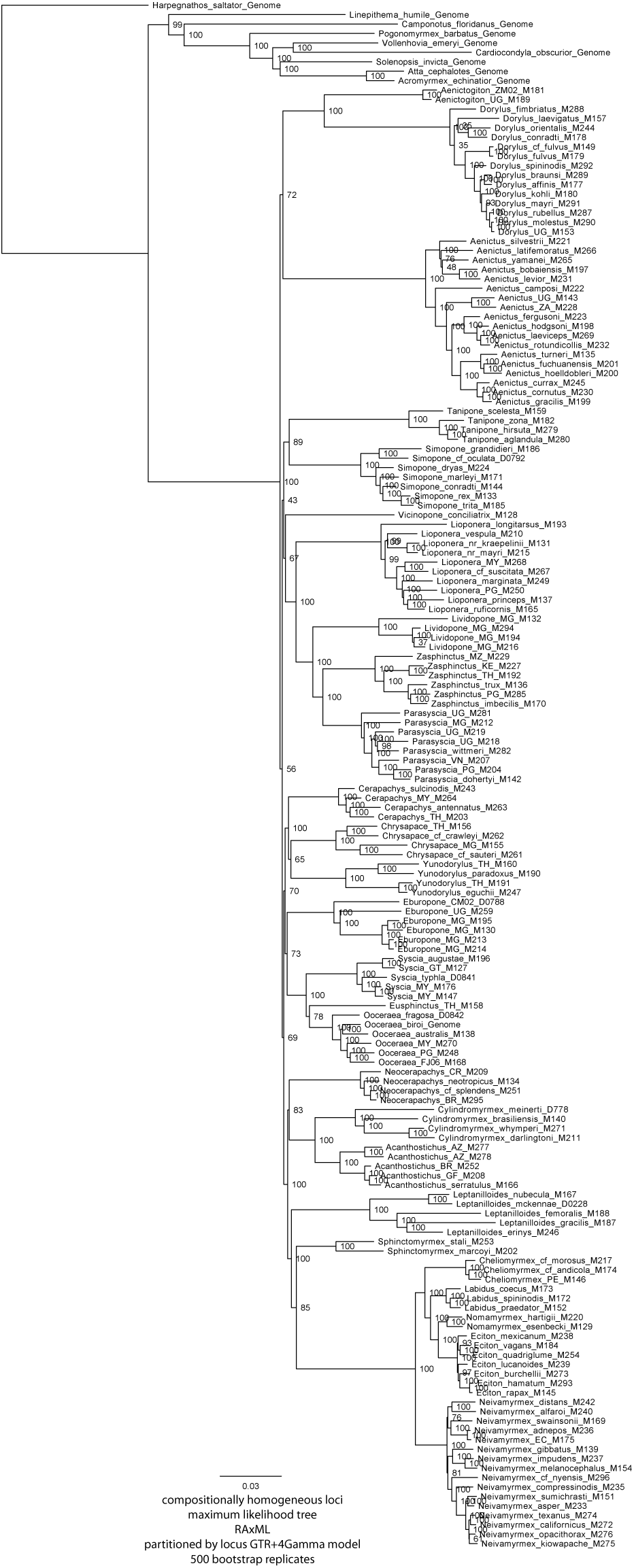
Maximum likelihood tree obtained from compositionally homogeneous data matrix partitioned by locus. Scale is in number of substitutions per site. Nodal support in percent bootstrap.

**Supplementary Figure 14:**
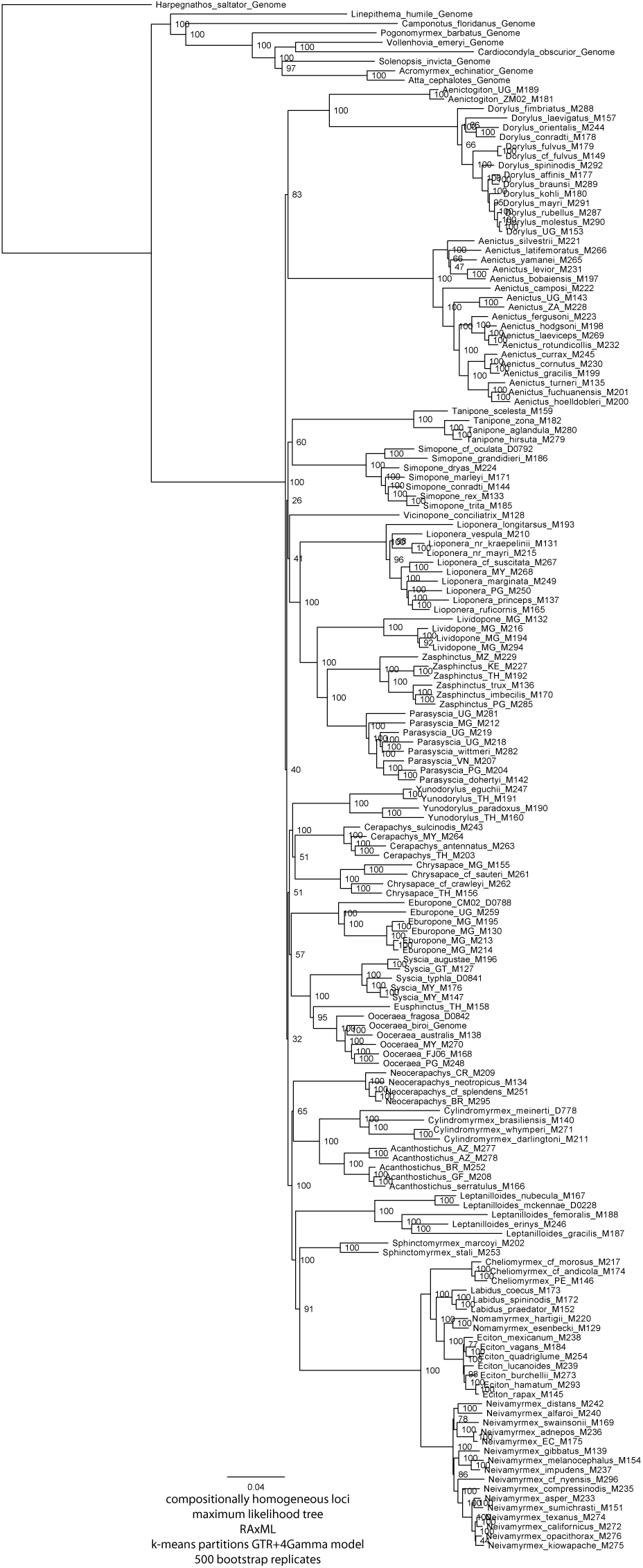
Maximum likelihood tree obtained from compositionally homogeneous data matrix under k-means partitions. Scale is in number of substitutions per site. Nodal support in percent bootstrap.

**Supplementary Figure 15:**
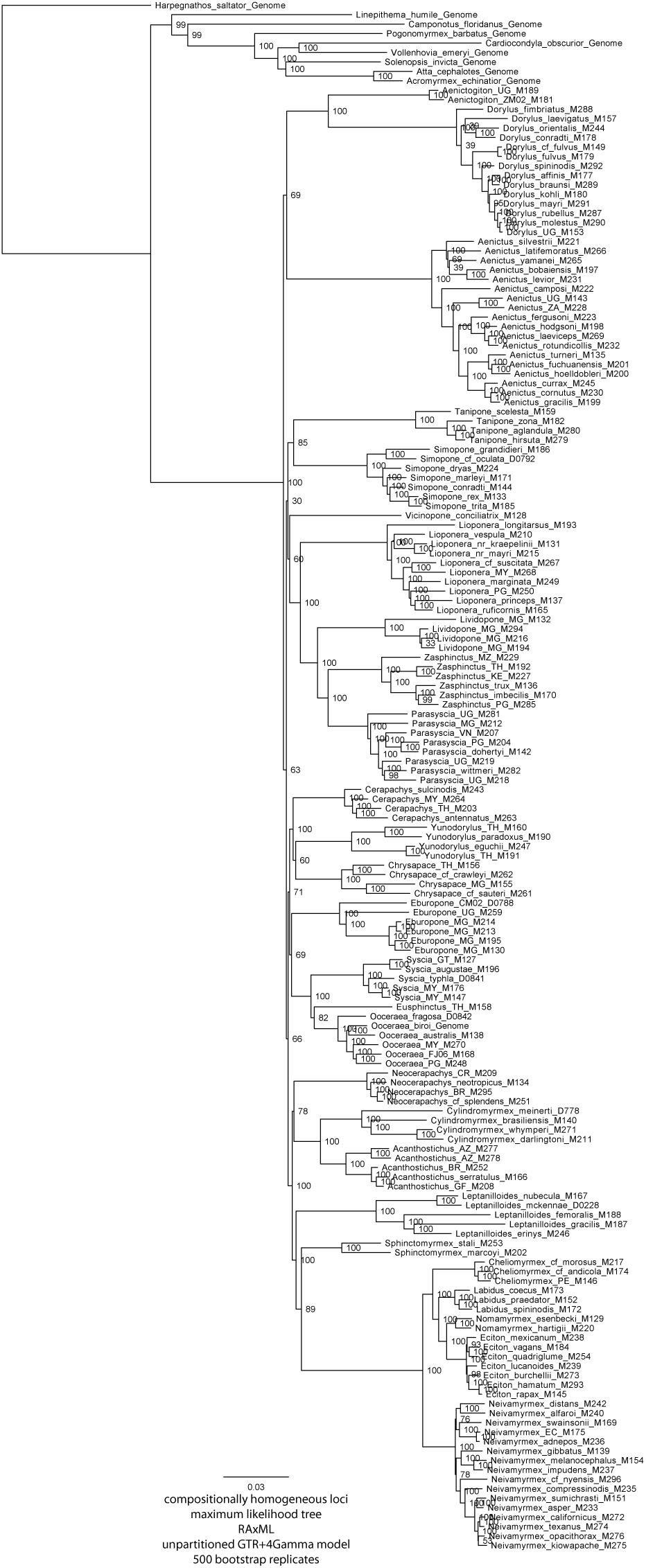
Maximum likelihood tree obtained from unpartitioned compositionally homogeneous data matrix. Scale is in number of substitutions per site. Nodal support in percent bootstrap.

**Supplementary Figure 16:**
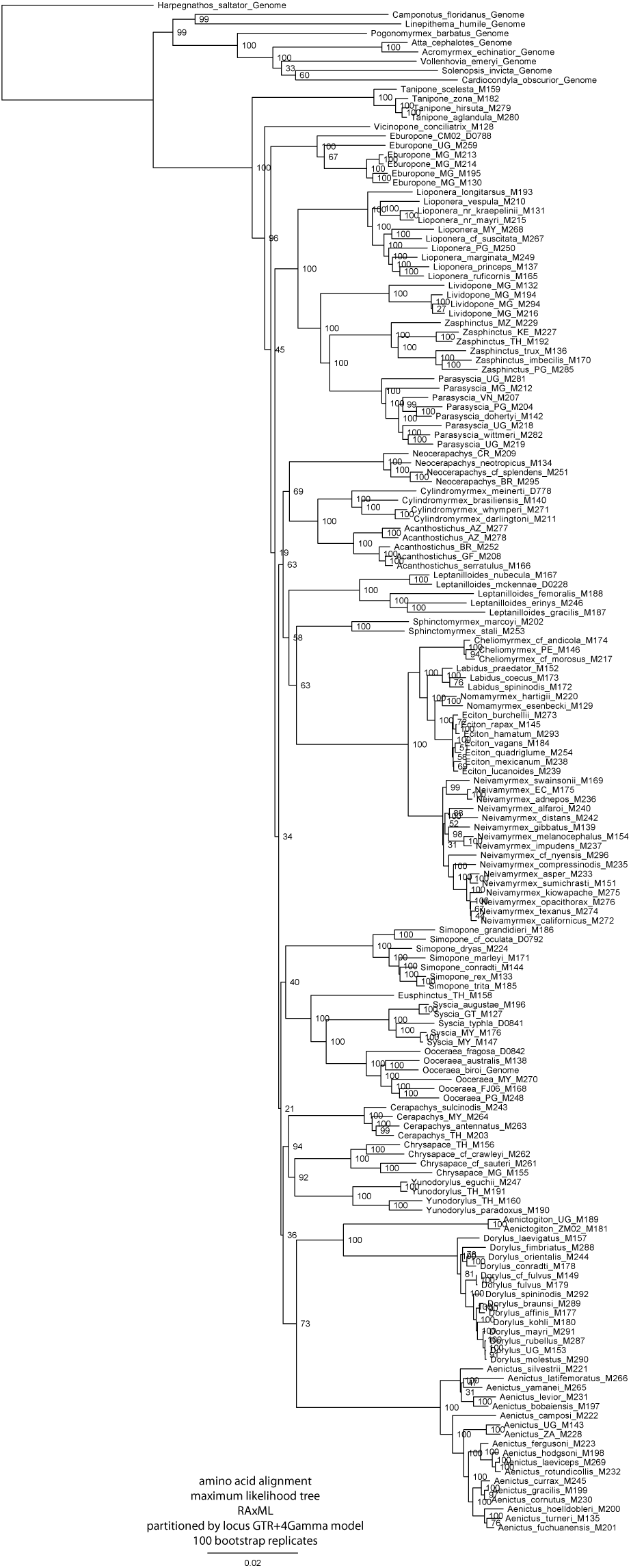
Maximum likelihood tree obtained from amino acid data matrix partitioned by locus. Scale is in number of substitutions per site. Nodal support in percent bootstrap.

**Supplementary Figure 17:**
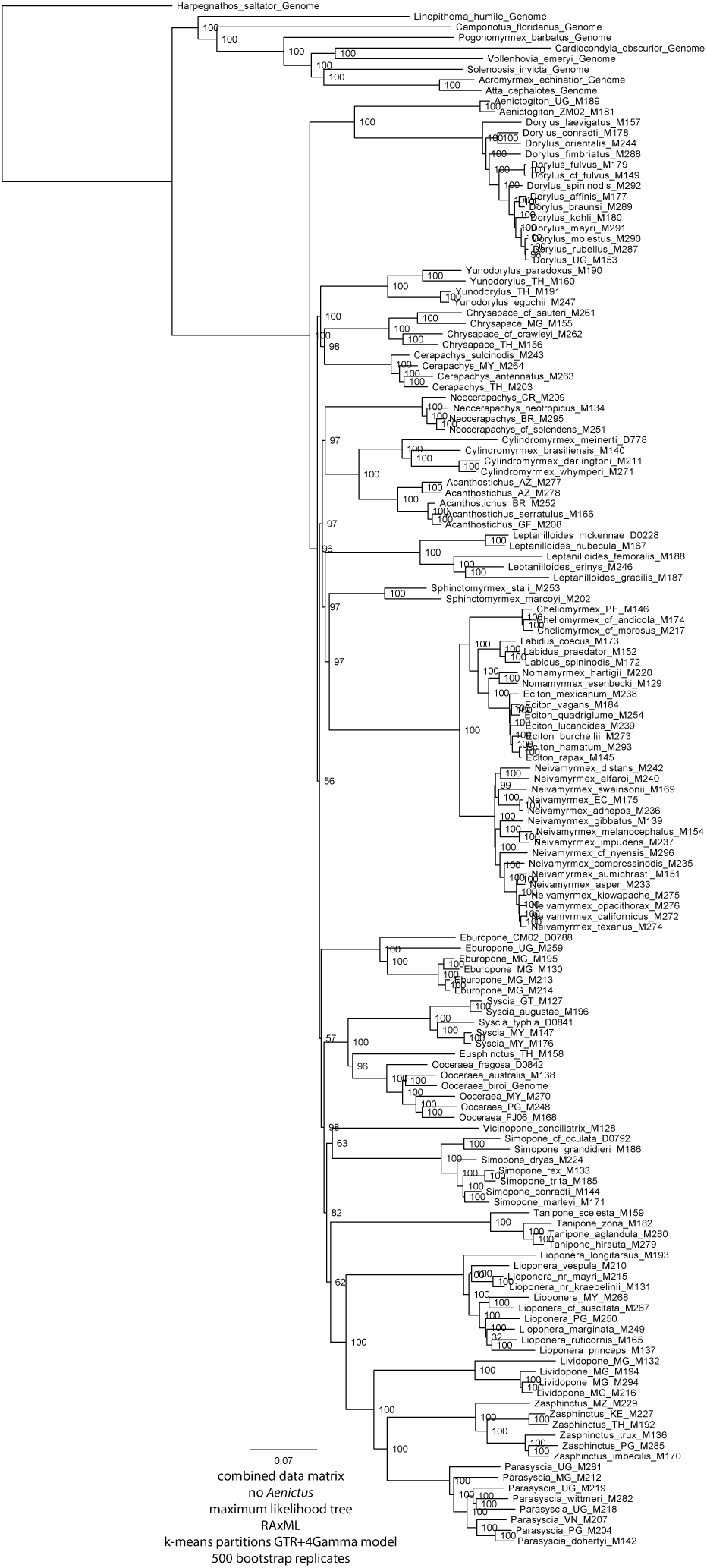
Maximum likelihood tree obtained under from complete data matrix with *Aenictus* removed under k-means partitioning. Scale is in number of substitutions per site. Nodal support in percent bootstrap.

**Supplementary Figure 18:**
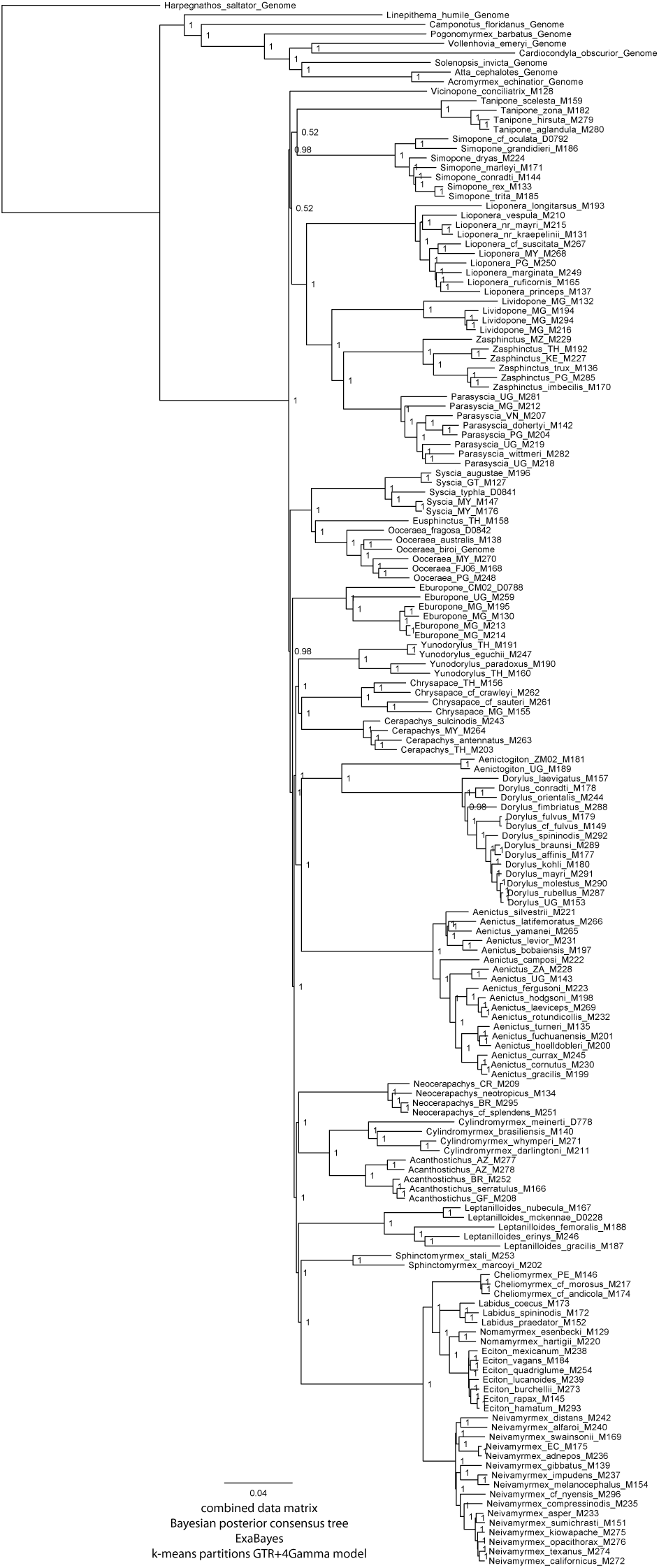
Bayesian posterior consensus tree obtained from complete data matrix under k-means partitioning. Scale is in number of substitutions per site. Nodal support in posterior probability.

**Supplementary Figure 19:**
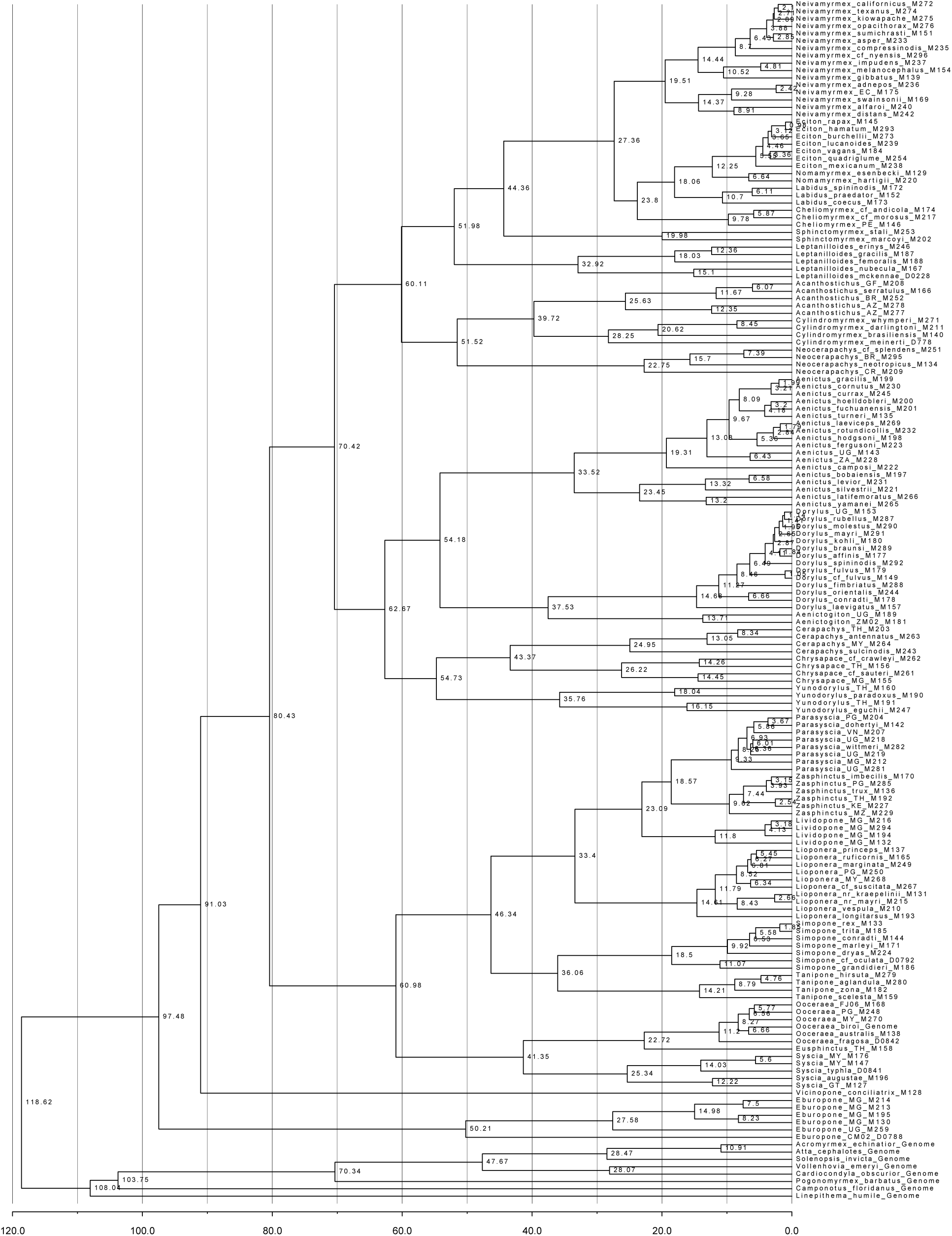
Summary tree with mean ages from 100 analyses under penalized likelihood in Chronos.

**Supplementary Figure 20:**
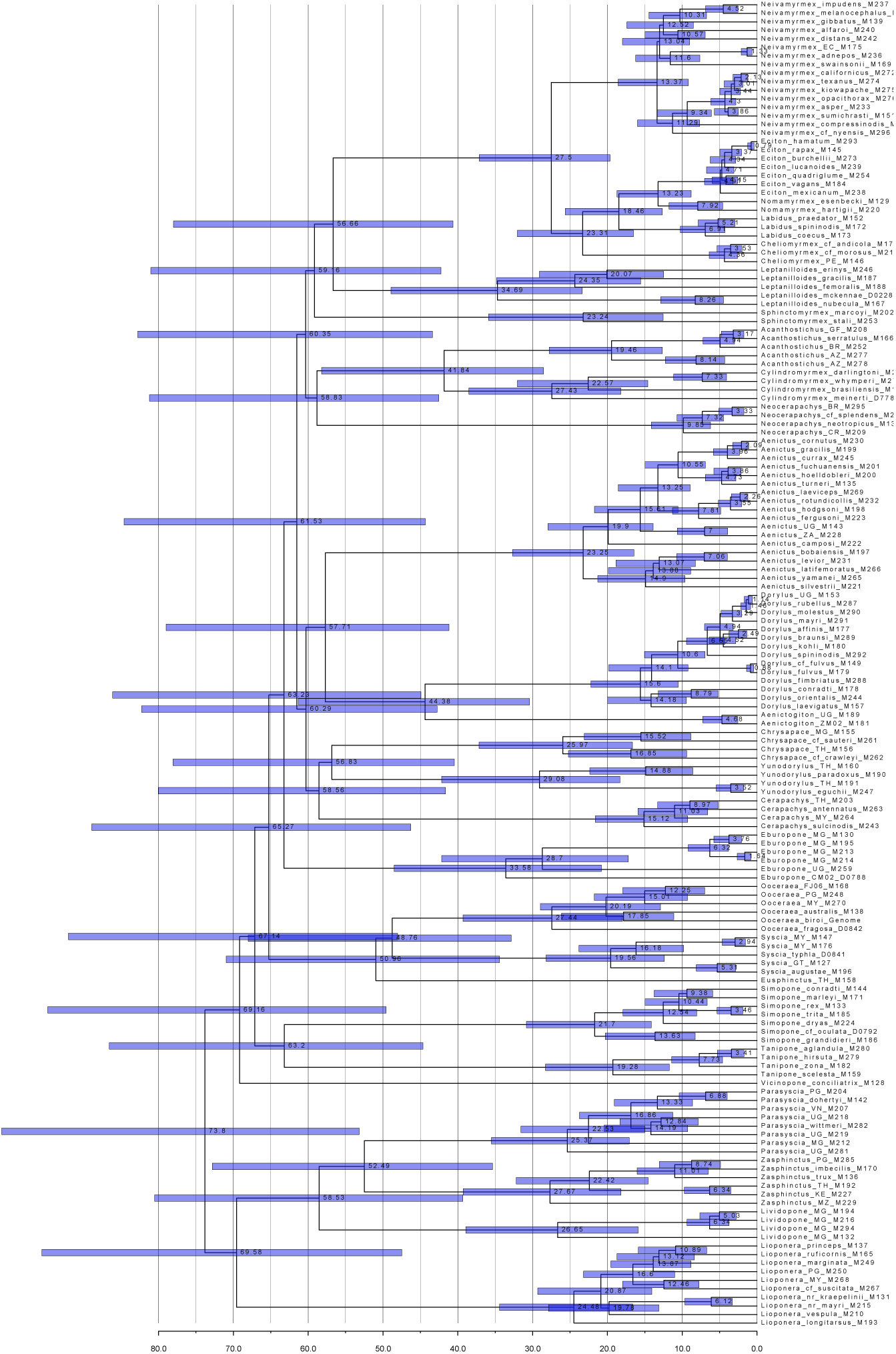
Mean ages and their 95% confidence intervals on the consensus BEAST tree inferred under fossilized birth-death process. All ages in Ma.

**Supplementary Figure 21:**
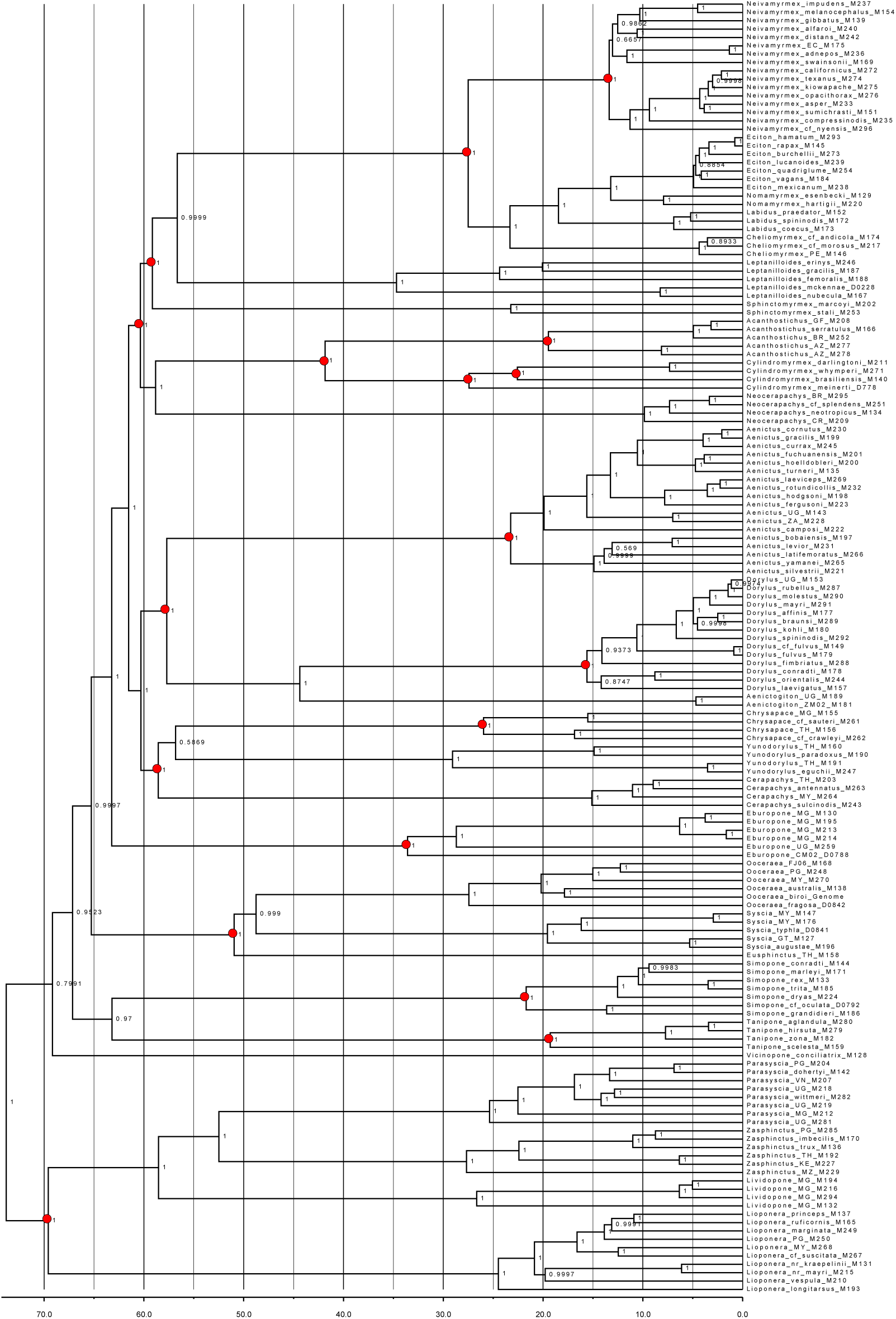
Posterior probabilities on the consensus BEAST tree inferred under fossilized birth-death process. Red dots reflect monophyly constraints used in the dating analysis. All ages in Ma.

**Supplementary Figure 22:**
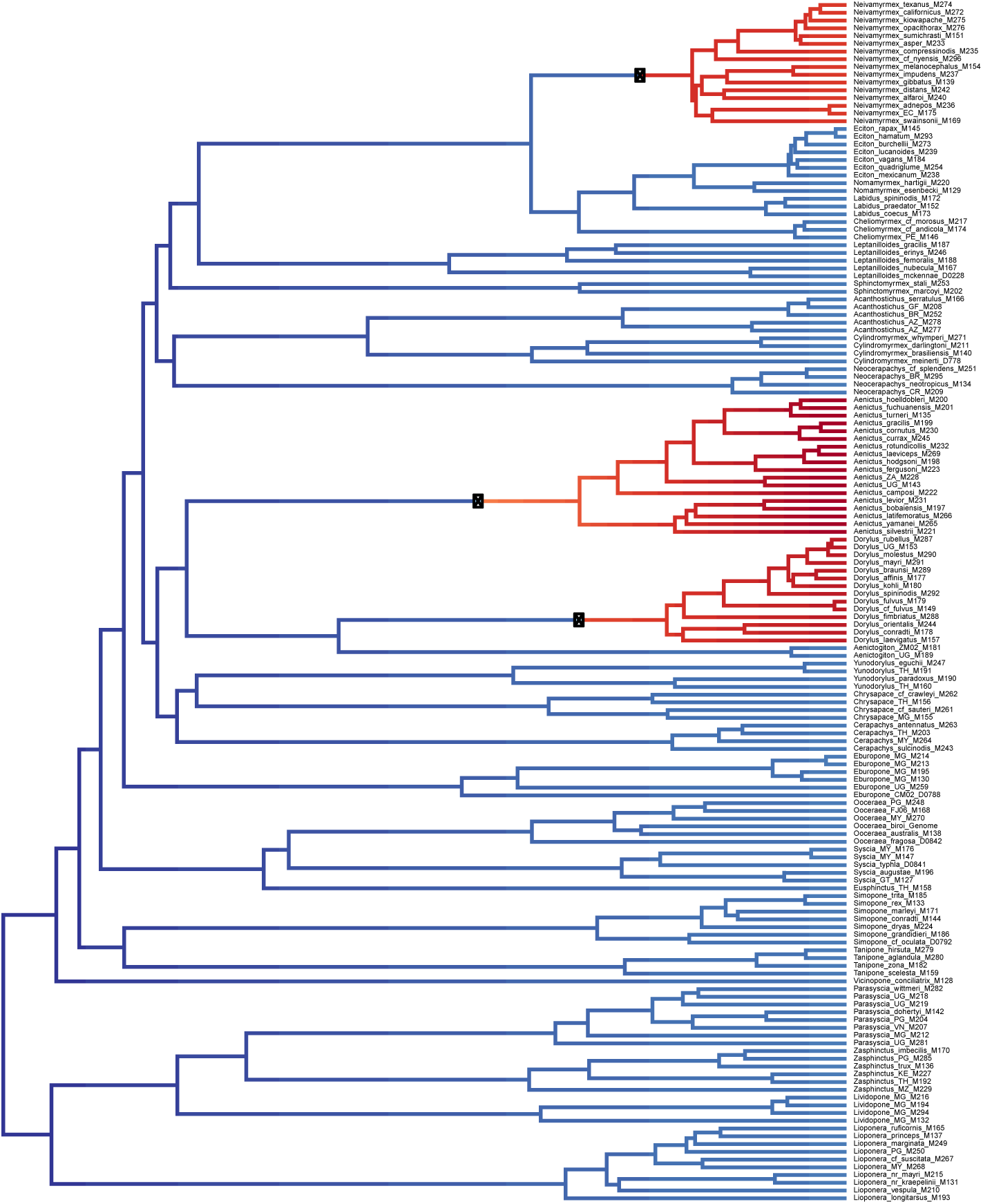
BAMM rate shift tree showing the overall best fit configuration. Red-filled circles signify placement of the shifts.

**Supplementary Figure 23:**
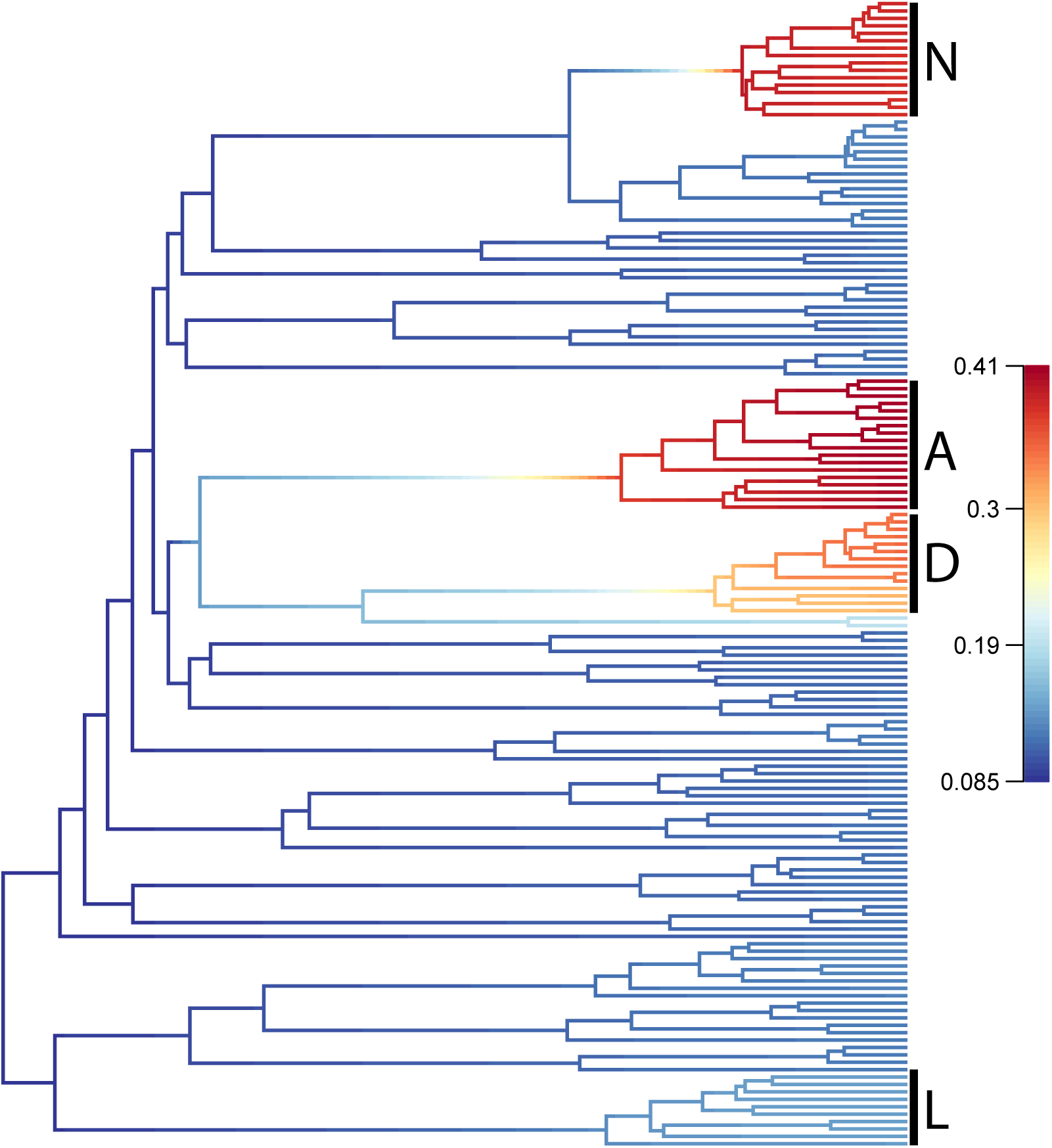
BAMM rate shift tree showing net diversification rates. A: *Aenictus*, D: *Dorylus*, L: *Lioponera*, N: *Neivamyrmex*.

**Supplementary Figure 24:**
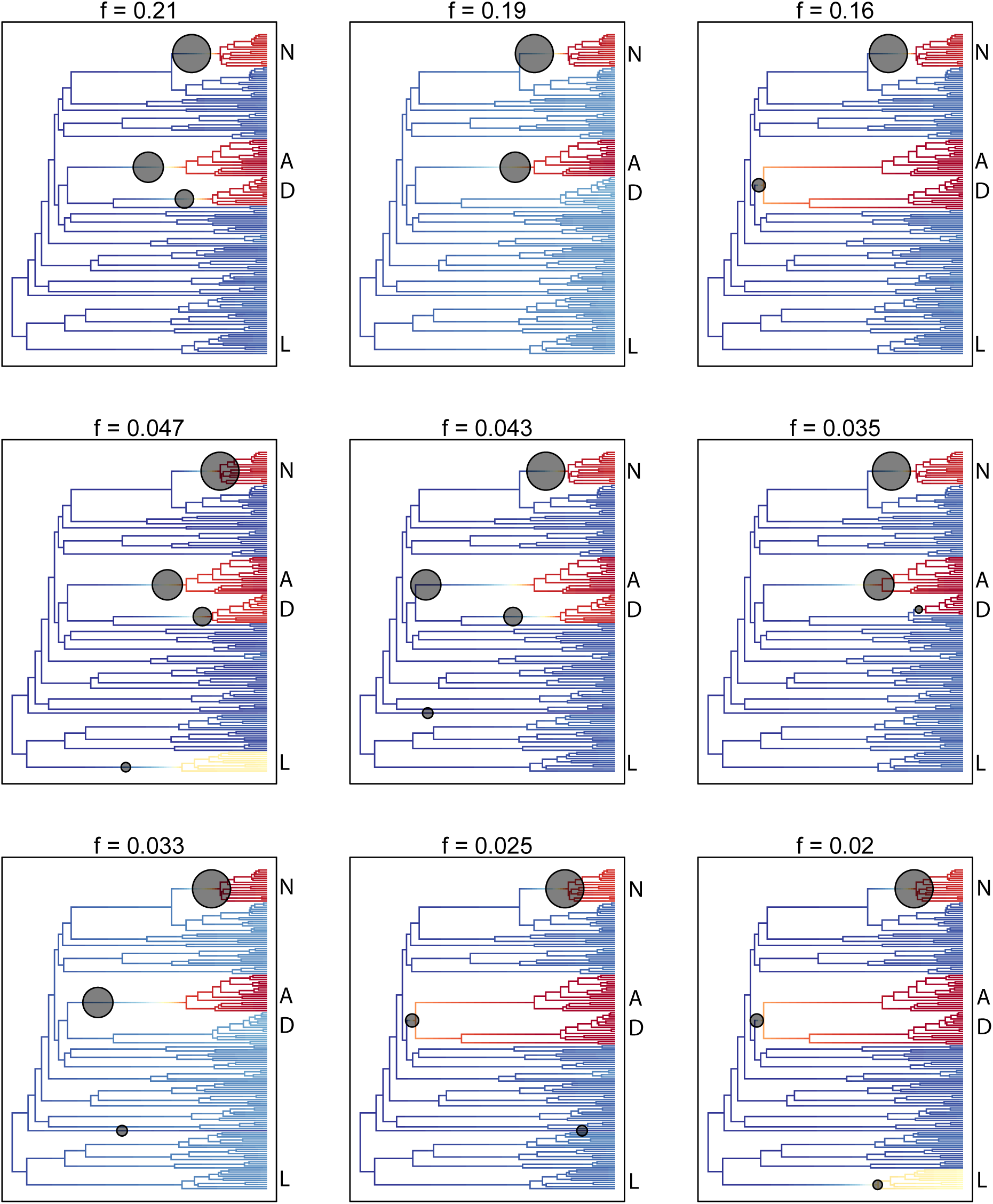
BAMM plot showing nine most common shift configurations in the credible set. The “f” number corresponds to the proportion of the posterior samples in which this configuration is present. A: *Aenictus*, D: *Dorylus*, L: *Lioponera*, N: *Neivamyrmex*.

**Supplementary Figure 25:**
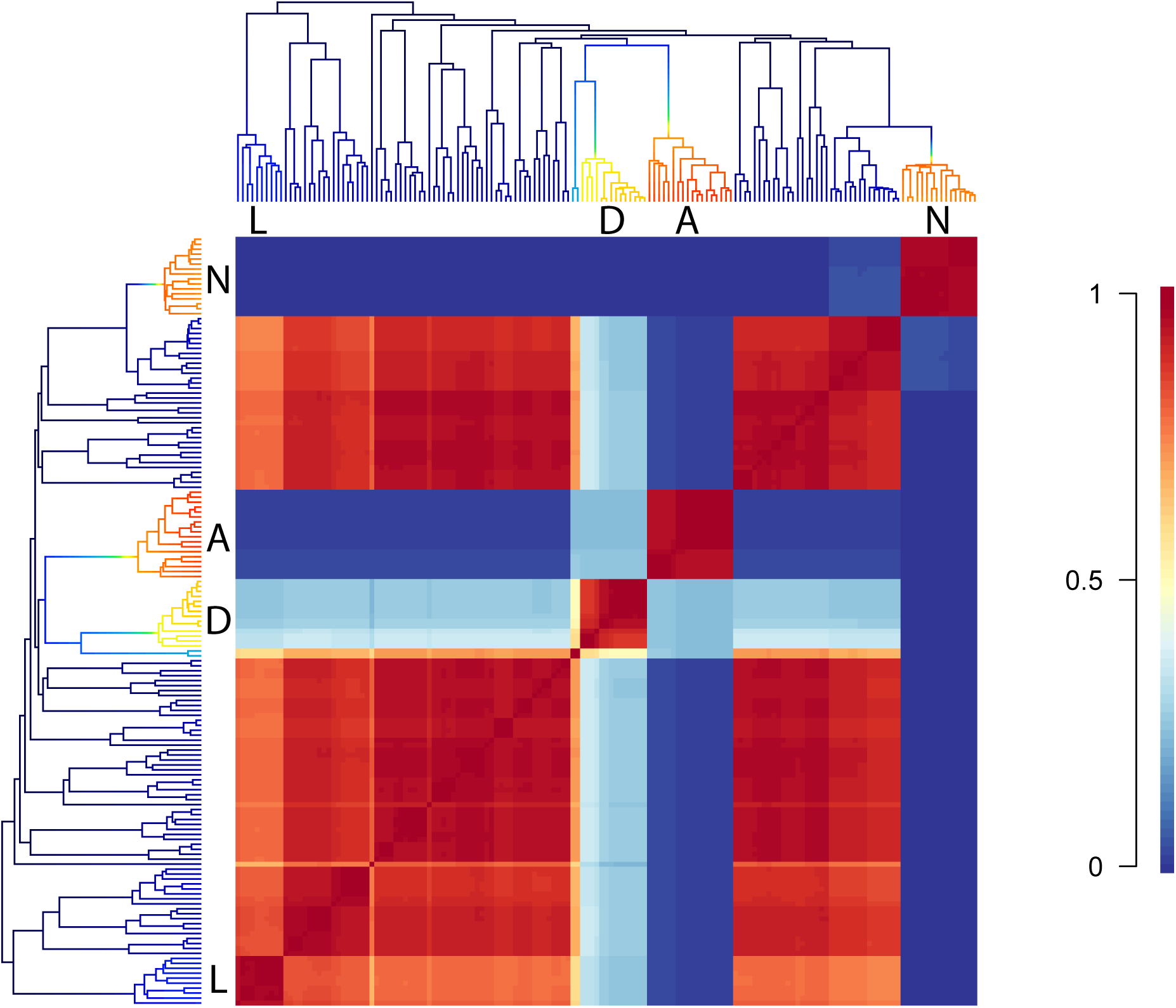
BAMM cohort plot. Blocks signify comparisons of shift regimes among species and clades, except across the diagonal which represents the comparison of a species to itself. A: *Aenictus*, D: *Dorylus*, L: *Lioponera*, N: *Neivamyrmex*.

**Supplementary Figure 26:**
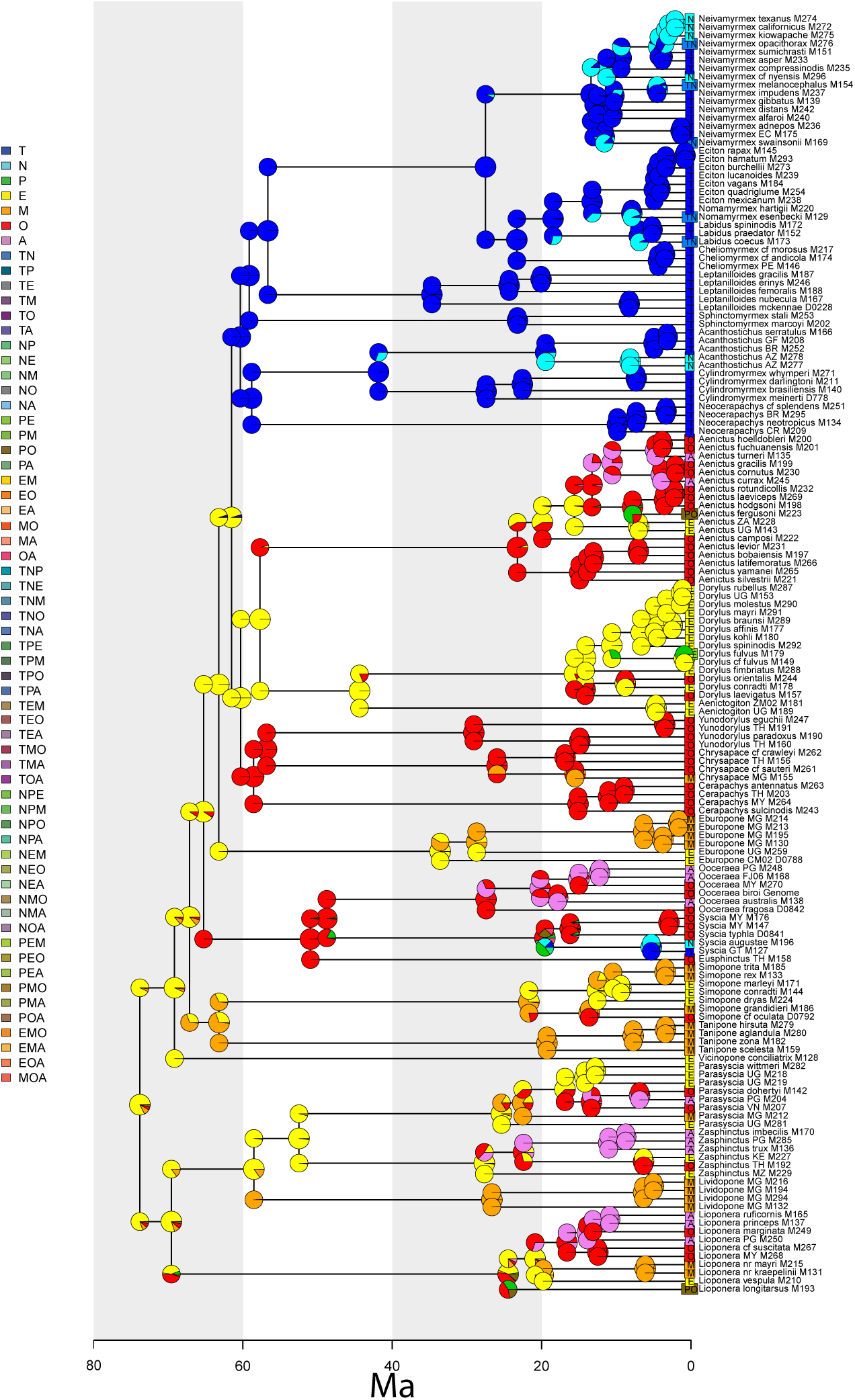
Relative likelihoods of ranges estimations from BioGeoBEARS under DEC+J, averaged over 100 posterior BEAST trees. Pie charts at the nodes correspond to ancestral state estimations and pie charts on the corners correspond to ranges immediately following speciation. The region names are abbreviated as follows: Neotropical (T), Nearctic (N), Palearctic (P), Afrotropical (E), Malagasy (M), Indomalayan (O), and Australasian (A). All ages in Ma.

**Supplementary Figure 27:**
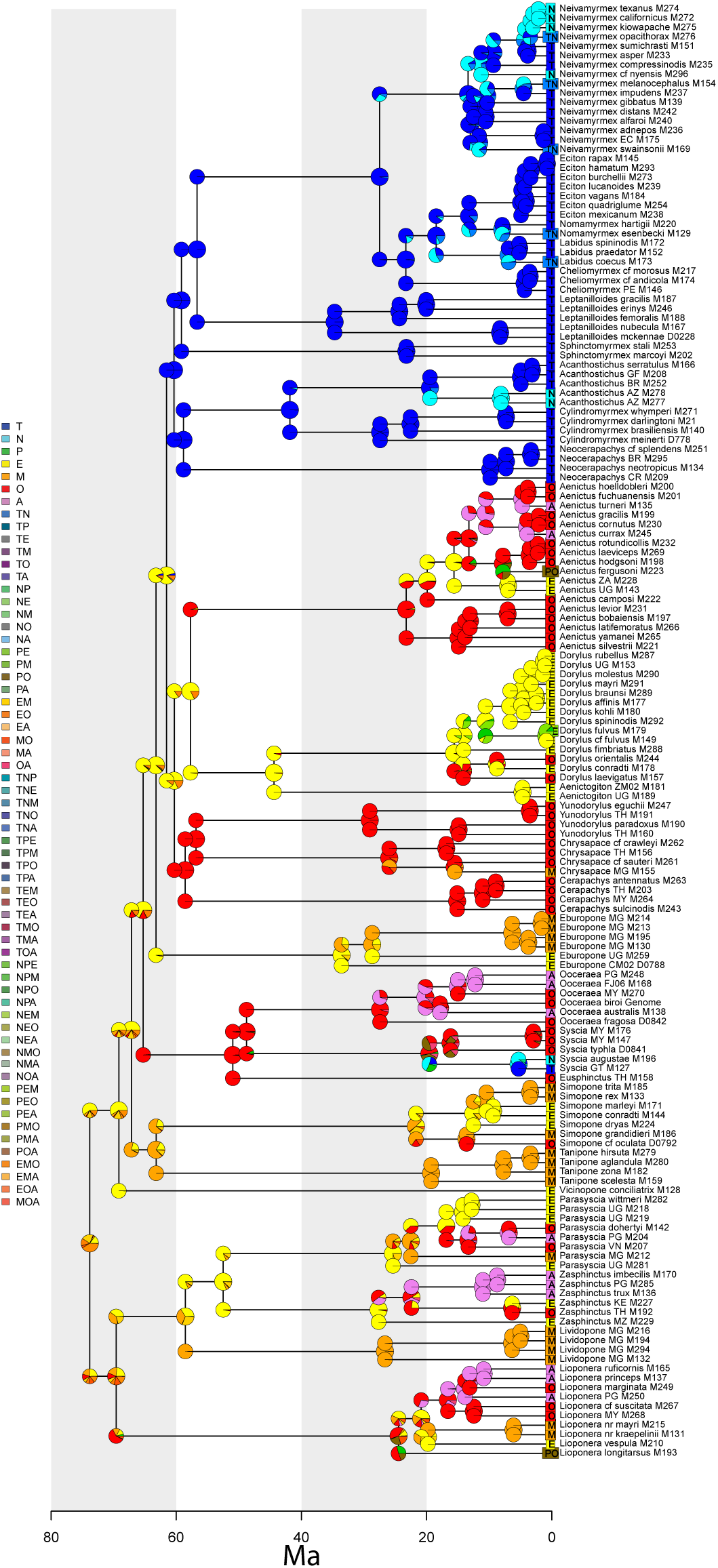
Relative likelihoods of ranges from BioGeoBEARS under DEC+J estimated on the BEAST consensus tree. Pie charts at the nodes correspond to ancestral state estimations and pie charts on the corners correspond to ranges immediately following speciation. The region names are abbreviated as follows: Neotropical (T), Nearctic (N), Palearctic (P), Afrotropical (E), Malagasy (M), Indomalayan (O), and Australasian (A). All ages in Ma.

**Supplementary Figure 28:**
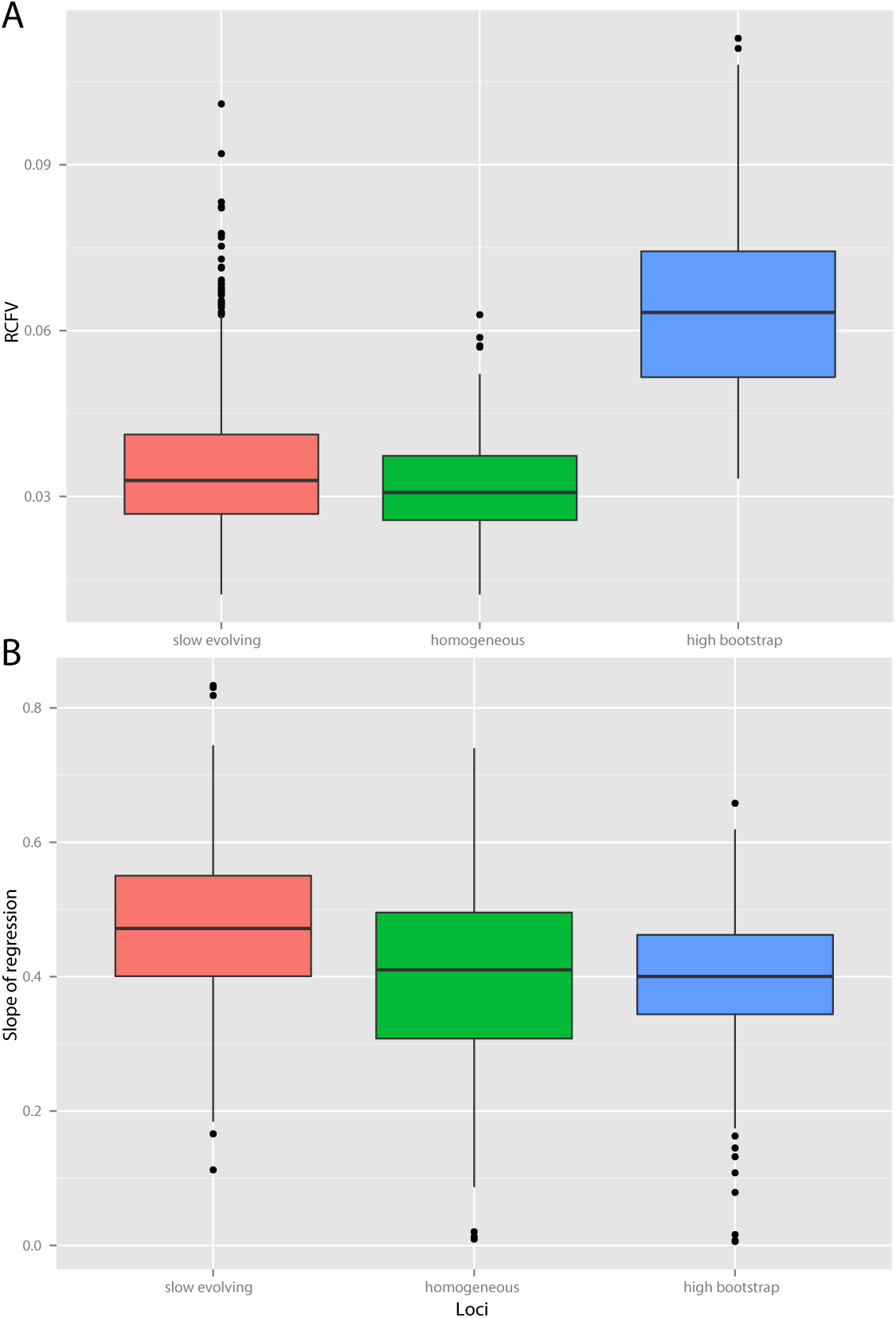
Box plots comparison of properties of slow-evolving, compositionally homogeneous, and “high signal” or high average bootstrap loci. A: Relative composition frequency variability (RCFV), B: Slope of regression of p-distances against distances on ML tree from a locus. Higher RCFV signifies more compositional heterogeneity and higher slope of regression signifies less potential for saturation.

**Supplementary Table 1:**
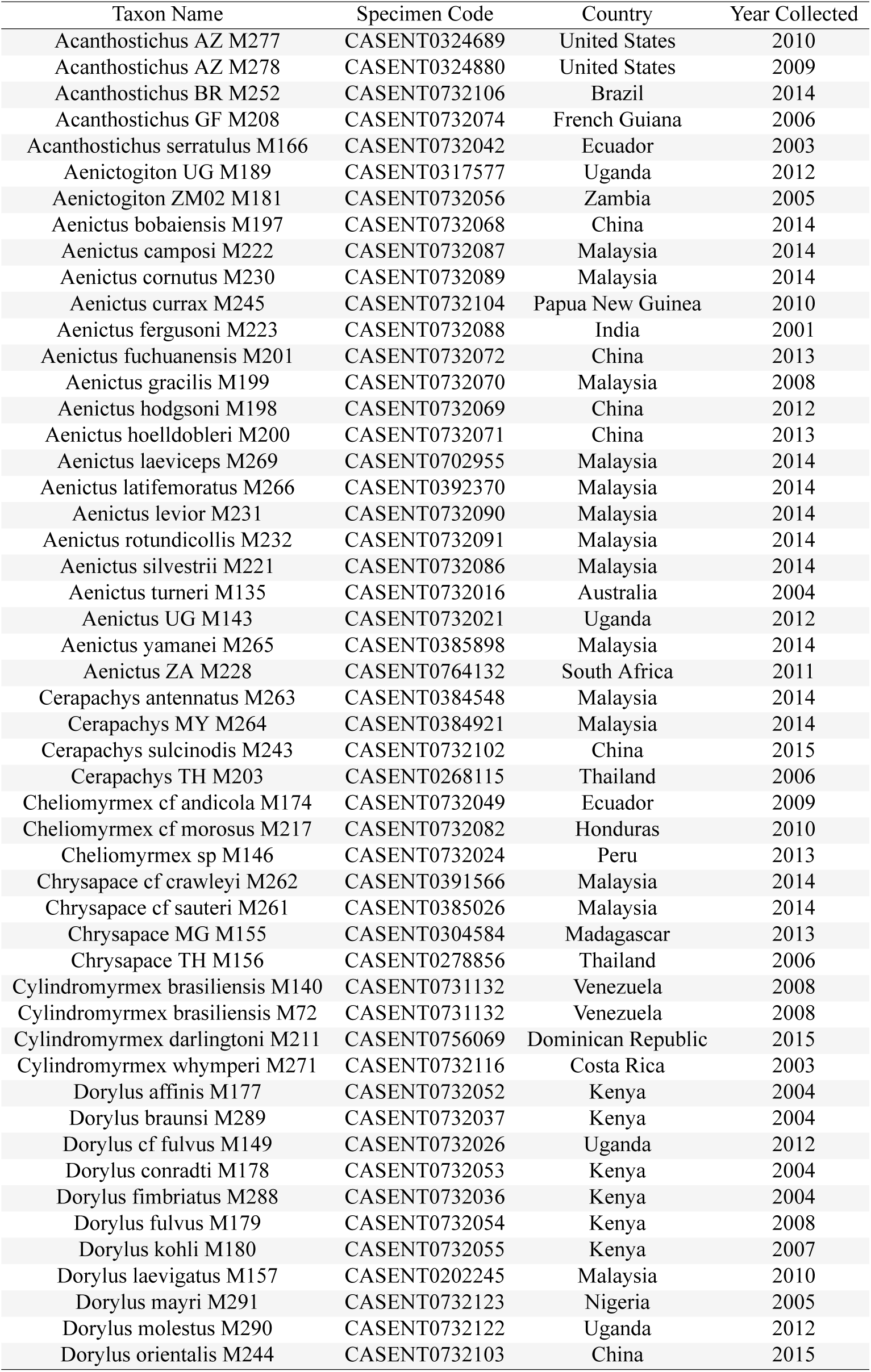
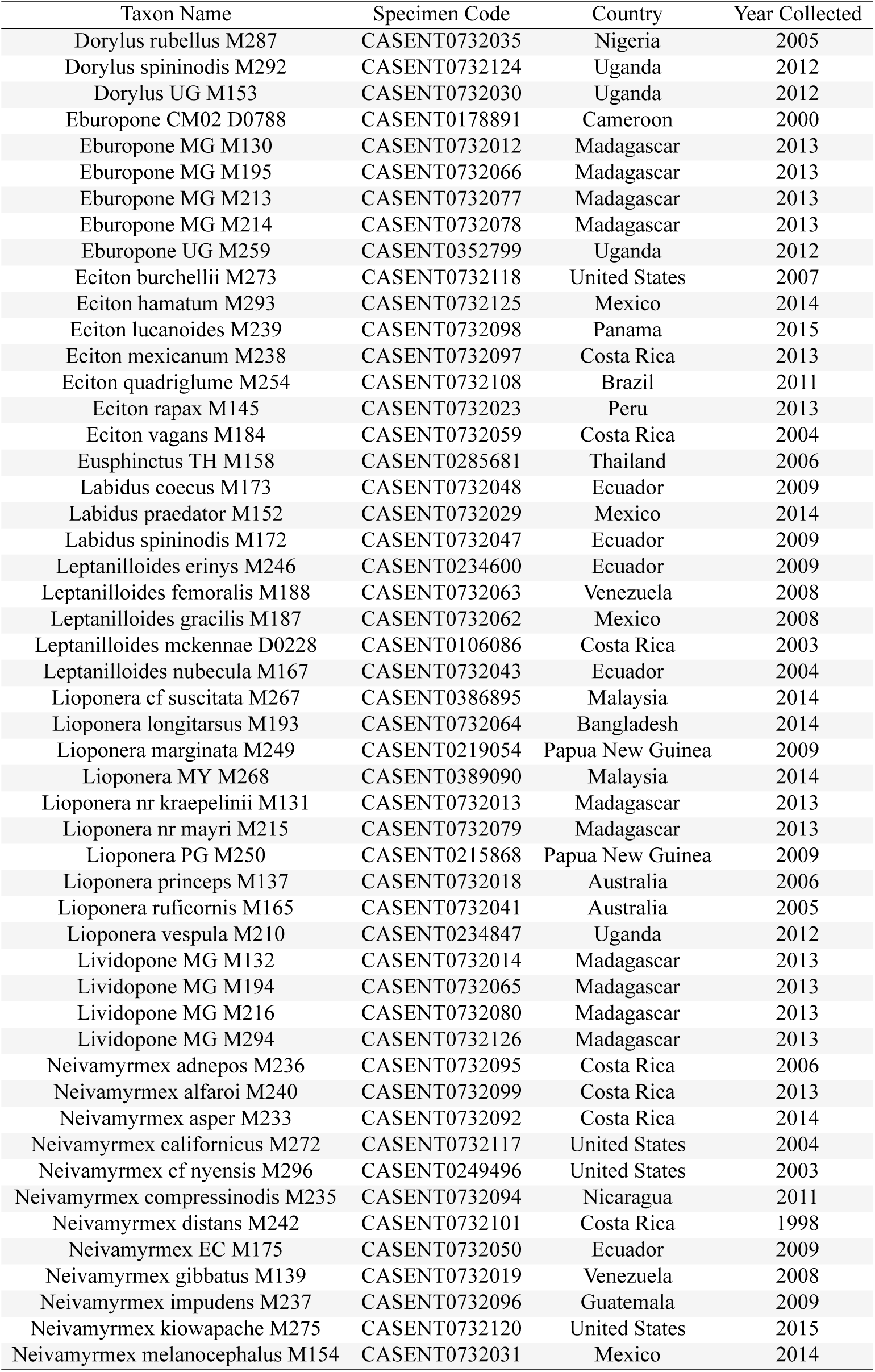
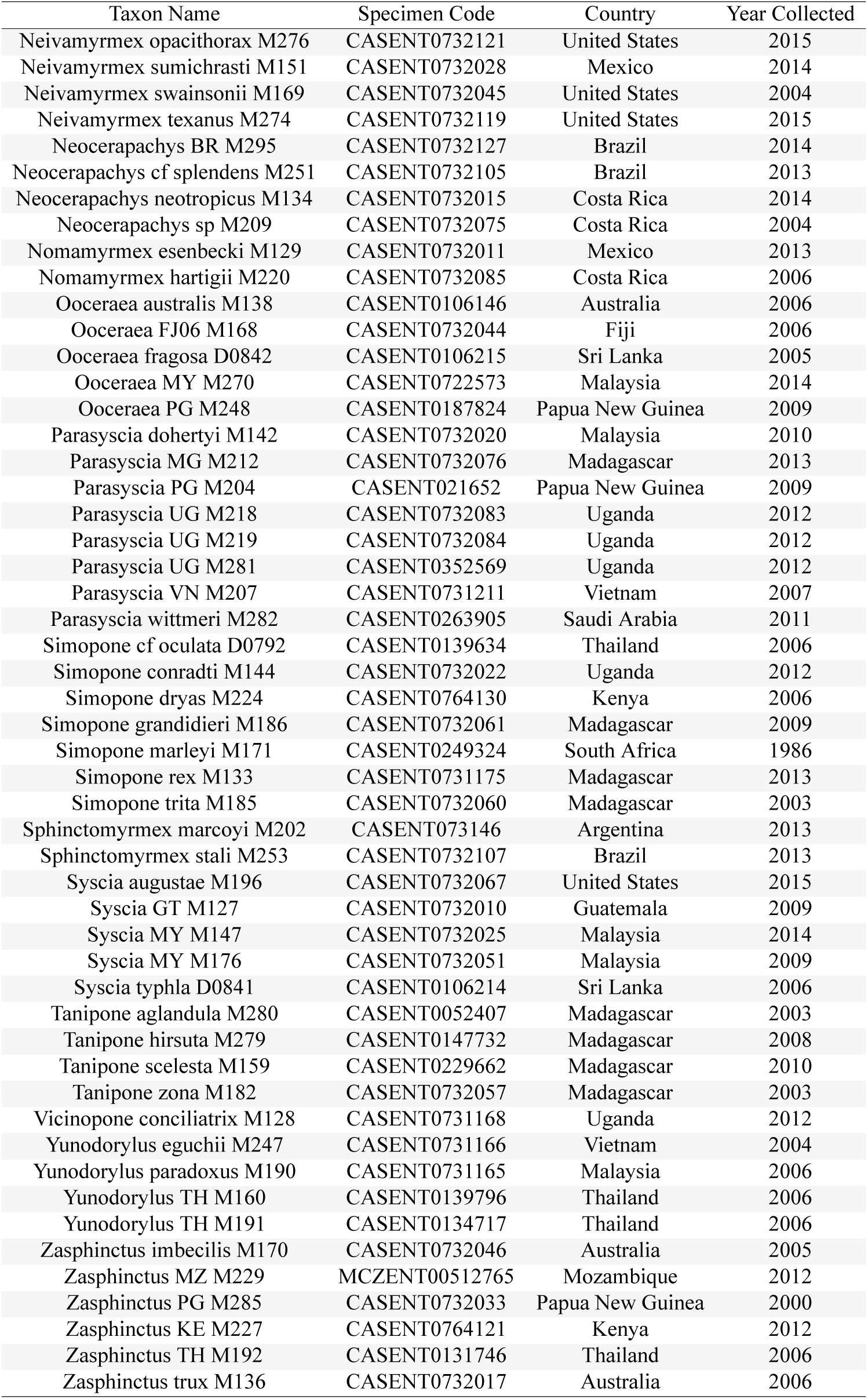
Voucher specimens used in this study. CASENT numbers correspond to records on AntWeb.org.

**Supplementary Table 2:** Statistics of data matrices used in this study.

| Alignment Name | Number of Taxa | Alignment Length | Percent Missing | Proportion of Parsimony Informative Sites | AT Content |
| --- | --- | --- | --- | --- | --- |
| Combined Data | 164 | 892,761 | 15.476 | 0.478 | 0.530 |
| "High-Signal" Loci | 164 | 177,947 | 17.534 | 0.565 | 0.578 |
| Slow-Evolving Loci | 164 | 178,662 | 10.245 | 0.348 | 0.486 |
| Homogeneous Loci | 164 | 97,849 | 10.177 | 0.369 | 0.488 |
| No *Aenictus* | 146 | 892,761 | 15.006 | 0.469 | 0.528 |
| BEAST Matrix | 155 | 44,079 | 16.489 | 0.466 | 0.531 |
| Amino Acids | 164 | 89,483 | 16.104 | 0.138 | - |

**Supplementary Table 3:**
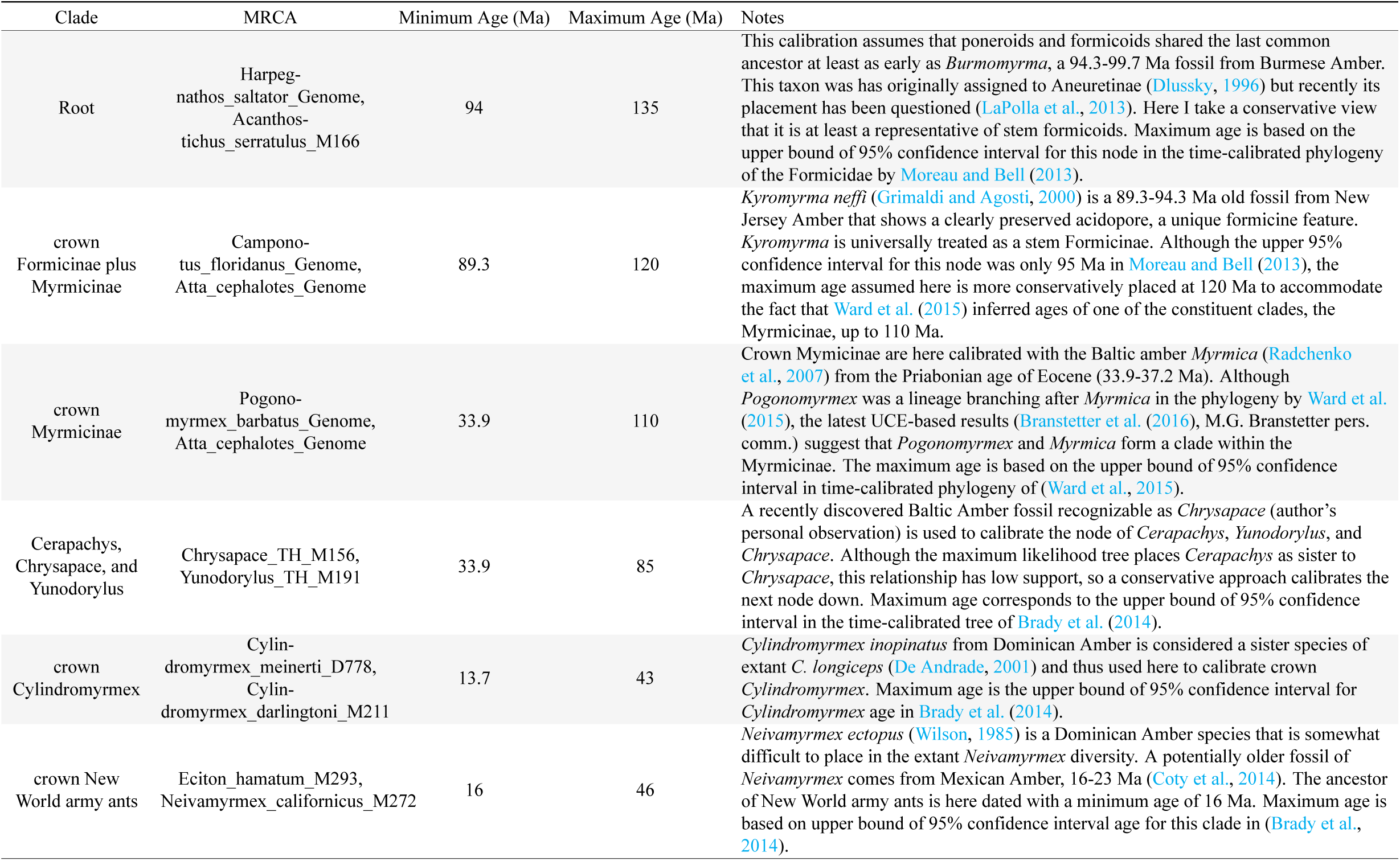
Calibration scheme used for penalized likelihood analyses in chronos. MRCA column refers to the most recent common ancestor of two tip names in the maximum likelihood tree obtained from slow-evolving loci matrix.

**Supplementary Table 4:**
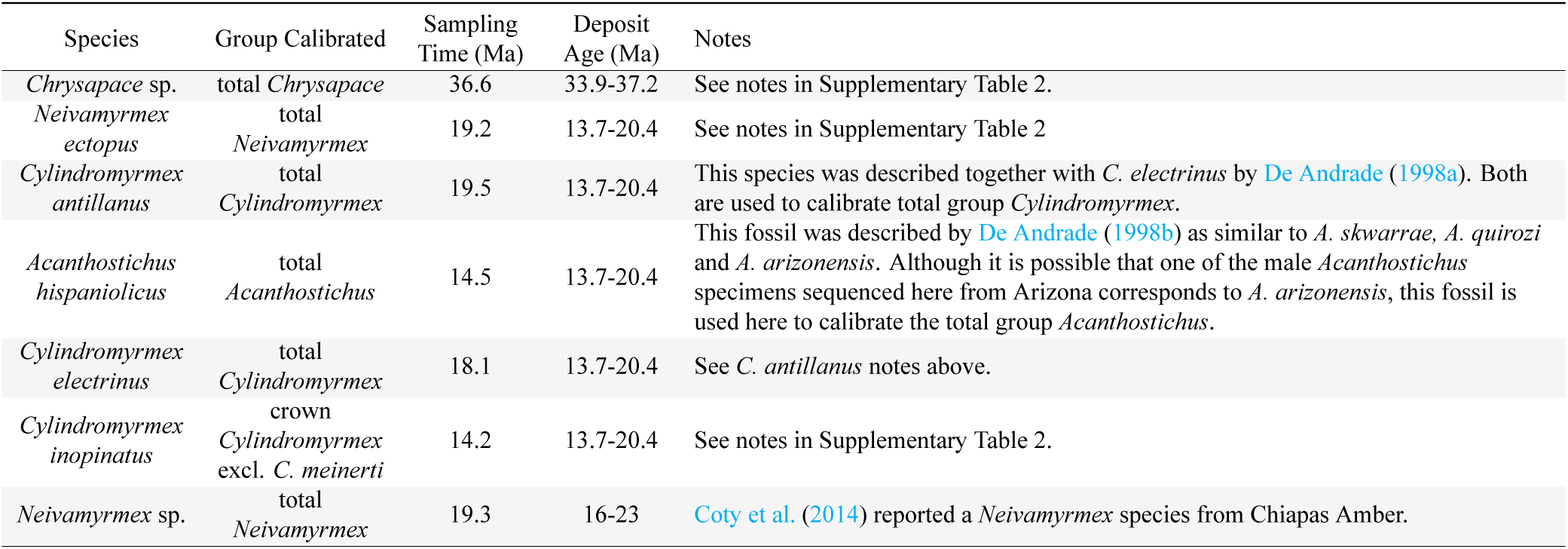
Calibration scheme used for fossilized birth-death process analyses in BEAST. Total group refers to fossil that could be placed in either stem or crown group.

| Species | Group Calibrated | Sampling Time (Ma) | Deposit Age (Ma) | Notes |
| --- | --- | --- | --- | --- |
| *Chrysapace* sp. | total *Chrysapace* | 36.6 | 33.9-37.2 | See notes in Supplementary Table 2. |
| *Neivamyrmex ectopus* | total<br>*Neivamyrmex* | 19.2 | 13.7-20.4 | See notes in Supplementary Table 2 |
| *Cylindromyrmex antillanus* | total<br>*Cylindromyrmex* | 19.5 | 13.7-20.4 | This species was described together with *C. electrinus* by De Andrade (1998a). Both are used to calibrate total group *Cylindromyrmex*. |
| *Acanthostichus hispaniolicus* | total<br>*Acanthostichus* | 14.5 | 13.7-20.4 | This fossil was described by De Andrade (1998b) as similar to *A. skwarrae*, *A. quirozi* and *A. arizonensis*. Although it is possible that one of the male *Acanthostichus* specimens sequenced here from Arizona corresponds to *A. arizonensis*, this fossil is used here to calibrate the total group *Acanthostichus*. |
| *Cylindromyrmex electrinus* | total<br>*Cylindromyrmex* | 18.1 | 13.7-20.4 | See *C. antillanus* notes above. |
| *Cylindromyrmex inopinatus* | crown<br>*Cylindromyrmex*<br>excl. *C. meinerti* | 14.2 | 13.7-20.4 | See notes in Supplementary Table 2. |
| *Neivamyrmex* sp. | total<br>*Neivamyrmex* | 19.3 | 16-23 | Coty et al. (2014) reported a *Neivamyrmex* species from Chiapas Amber. |

